# SPECC1L binds MYPT1/PP1β and can regulate its distribution between microtubules and filamentous actin

**DOI:** 10.1101/2022.09.09.507377

**Authors:** V Mehta, N Decan, A Gaudreau-Lapierre, JW Copeland, L Trinkle-Mulcahy

## Abstract

The subcellular localization, activity and substrate specificity of the serine/threonine protein phosphatase 1 catalytic subunit (PP1 _cat_) is mediated through its dynamic association with regulatory subunits in holoenzyme complexes. While some functional overlap is observed for the three human PP1_cat_ isoforms, they also show distinct targeting based on relative preferences for specific regulatory subunits. A well-known example is the preferential association of MYPT1 with PP1β in the myosin phosphatase complex. In smooth muscle, MYPT1/ PP1β counteracts the muscle contraction induced by phosphorylation of the light chains of myosin by the myosin light chain kinase. This phosphatase complex is also found in non-muscle cells, where it is targeted to both myosin and non-myosin substrates and contributes to regulation of the balance of cytoskeletal structure and motility during cell migration and division. Although it remains unclear how MYPT1/PP1β traffics between microtubule- and actin-associated substrates, our identification of the microtubule- and actin-binding protein SPECC1L in both the PP1β and MYPT1 interactomes suggested that it may be the missing link. Validation of their association, together with the strong overlap that we observed for the SPECC1L and MYPT1 interactomes, suggested that they exist in a stable complex in the cell. We further showed that SPECC1L binds MYPT1 directly, and that it can impact the balance of the distribution of the MYPT1/ PP1β complex between the microtubule and filamentous actin networks.

## INTRODUCTION

Reversible protein phosphorylation is the most common post-translational modification, acting as a molecular switch that can modulate protein conformation and/or protein-protein interactions. This in turn leads to alterations in enzymatic activity, subcellular localization, turnover of targets or signaling by other PTMs. Phospho-regulation plays a role in most cellular processes, including signaling, metabolism, migration and cell cycle progression, and is a key therapeutic target in diseases in which these processes are deregulated. The predominant phosphorylated amino acid is Serine (Ser), which accounts for >80% of phosphorylation events [1]. Threonine (Thr) and Tyrosine (Tyr) account for the bulk of the remaining phospho-sites, with phosphoryation also demonstrated to a lesser extent on other amino acid residues [2]. Protein Phosphatase 1 (PP1) catalytic subunit (PP1_cat_) is ubiquitiously expressed in eukaryotic cells and estimated to account for up to 70% of Ser/Thr dephosphorylation events [3]. Mammalian PP1_cat_ is found primarily as three isoforms (α, β/δ, γ) that are encoded by three distinct genes [4]. These isoforms are >89% identical in amino acid sequence, with minor variations primarily at their NH_2_ and COOH termini [5]. Loss of function and biochemical studies of individual PP1 isoforms in eukaryotic organisms suggest some level of compensation or overlapping function, while also highlighting distinct phenotypes associated with the disruption of a single gene [4].

In the cell, PP1 is regulated and achieves its substrate specificity through the association of the catalytic subunit with a range of regulatory or “targeting” subunits [6]. This results in the combinatorial generation of a large and diverse group of dimeric PP1 holoenzyme complexes, each with its own subset of substrates and mechanism(s) of regulation. To date, >200 confirmed PP1 interacting proteins have been identified using a range of proteomic, bioinformatic, yeast two-hybrid and biochemical approaches [7]–[13]. The majority of known PP1 regulatory proteins contain an RVxF docking motif that mediates association with a hydrophobic channel in PP1_cat_ [14]. Several contain additional PP1-binding sequences, such as SILK and MyPhoNE motifs, that enhance binding and contribute to isoform preference (see [6] for review). There is also a subset of inhibitory PP1 interactors that do not contain a typical RVxF docking motif (e.g. Inh2, SDS22) and can associate with PP1 bound to another regulatory subunit in trimeric complexes [6].

Our interactome screens comparing the three human PP1 phosphatase isoforms have identified and characterized numerous novel proteins not previously defined as phosphatase complex members [7], [9], [15]. Of particular interest was the identification of the related Sperm antigen with calponin homology and coiled-coil domains 1 (SPECC1) and SPECC1L proteins, which were found to preferentially associate with PP1β yet possess no obvious PP1 binding motifs. That suggested that their interaction with PP1 is indirect, which was confirmed by their appearance in our MYPT1 interactome screens. MYPT1 is a regulatory subunit that binds preferentially to PP1β to generate the myosin phosphatase (MP) complex. Although most abundantly expressed in smooth muscle cells [15]–[17], it is found in many cell types, and targeted disruption of the Mypt1 gene in mice is embryonic lethal [18]–[20]. In addition to its canonical role in regulating muscle contraction via dephosphorylation of myosin light chains, the MP complex plays a key role in the regulation of actomyosin in non-muscle cells, affecting cell migration and adhesion [21]. Several non-myosin Mypt1/PP1β substrates have also been identified (for review see [22]). They include polo-like kinase 1 (PLK1), with which MYPT1 associates at centrosomes to contribute to mitotic regulation [23] and Histone Deacetylase 6 (HDAC6), which plays an important role in microtubule deacetylation [24].

Although not studied to the same extent, depletion of SPECC1L in cells led to defects in cytoskeletal organization and cell migration[25], and SPECC1L mutations have been linked to developmental facial morphogenesis disorders that result in congenital malformations [25]–[29]. The protein is found predominantly in the cytoplasm and has been shown to accumulate at both microtubule (MT) and filamentous actin structures throughout the cell cycle [25][30]. It is not yet clear how this distribution is regulated, as SPECC1L does not contain any obvious microtubule binding domains. A single calponin homology (CH) domain [31] at its C-terminus may facilitate actin binding, however two tandem CH domains are normally required to bind actin.

We confirmed that SPECC1L forms a stable complex with MYPT1 in non-muscle cells, that the binding is direct, and that it is mediated by their respective C-termini. Consistent with this, quantitative proteomic experiments demonstrated significant overlap of their interaction profiles, identifying proteins involved in the regulation of cell contractility, actin organization, microtubule stability, junction turnover, cytokinesis, adhesion and migration. In addition to mapping the regions of SPECC1L that mediate association with MYPT1, MTs and actin, we also demonstrated its ability to modulate the distribution of MYPT1/PP1β between these two cytoskeletal networks.

## RESULTS

### SPECC1L/1 associate with the myosin phosphatase complex

As part of our ongoing analysis of PP1 holoenzyme complexes, we used our quantitative SILAC (Stable Isotope Labeling by Amino Acids in Culture) affinity purification/mass spectrometry (AP/MS) approach [32] to map the interactome of MYPT1 in U2OS cells (Fig. 1a) and compared the overlapping enriched proteins in two independent datasets (Fig. 1b; Supp Data File 1). As expected, we saw strong enrichment of both PP1β and the myosin regulatory light chain, along with numerous proteins related to motility and cytoskeletal organization. We also observed strong enrichment of SPECC1L. To further validate their interaction, we demonstrated co-precipitation of endogenous SPECC1L with GFP-tagged MYPT1 from U2OS cell lysates (Fig. 1c), and the reciprocal co-precipitation of both MYPT1 and PP1β with GFP-tagged SPECC1L (Fig. 1d). In a complementary BioID approach [33], we demonstrated biotin-based proximity labeling of MYPT1 in cells expressing SPECC1L fused to the biotin ligase BirA* (Fig. S1a). Enrichment of the related family member SPECC1 was also detected in one of the MYPT1 datasets (Supp Data File 1), and its association further validated by both co-IP of MYPT1 with GFP-tagged SPECC1 (Fig. S1e) and by proximity labeling of MYPT1 in cells expressing BirA*-tagged SPECC1 (Fig. S1b). While we were carrying out these experiments, a SPECC1L-MYPT1 association was also annotated in a large-scale screen for phosphatase interactors[13], although it was not further explored in that study.

**Figure 1.**
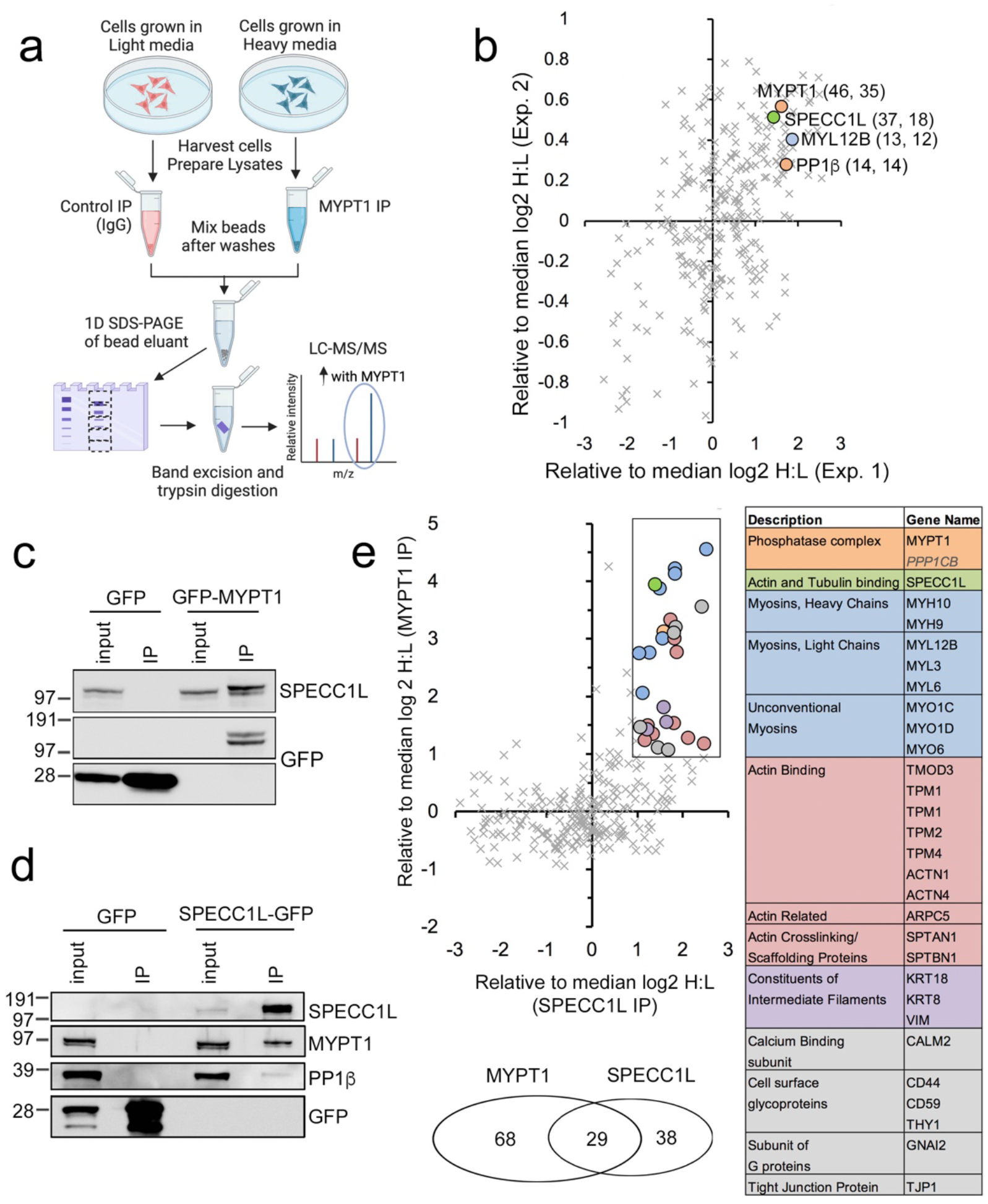
SPECC1L associates with the myosin phosphatase complex and shows an overlapping interactome enriched in cytoskeletal factors. a. Design of the SILAC AP/MS experiment used to map the interactome of endogenous MYPT1. b. graph showing factors enriched in 2 replicates of the AP/MS experiment (upper right quadrant), plotted as the displacement from the median log2 H:L ratio in both. For the highlighted proteins, the number of peptides detected in each experiment is noted in parentheses. c. GFP-MYPT1 immunoprecipitated from U2OS cells co-purifies endogenous SPECC1L. d. SPECC1L-GFP immunoprecipitated from U2OS^SPECC1L-EGFP^ cells co-purifies MYPT1 and PP1β. e. Factors enriched >2-fold in both MYPT1 and SPECC1L SILAC AP/MS experiments are highlighted on the graph (upper right quadrant) and colour-coded table. PP1β (in italics) was just below the 2-fold enrichment threshold in the SPECC1L experiment. The Venn diagram shows the number of factors that overlap. The full datasets are provided in Supplemental Data File 1.

We next mapped the interactome of SPECC1L in U2OS cells, to compare its overlap with that of MYPT1. As none of the available commercial antibodies were suitable for immunoprecipitation, we established a U2OS cell line stably overexpressing GFP-tagged SPECC1L at endogenous levels (Fig. S3b) and confirmed that its subcellular localization matched that of the endogenous protein (Fig. 4a-b). Figure 1e shows a comparison of the SPECC1L-GFP and MYPT1 interactomes. As highlighted in the Venn diagram, the high confidence hits (top right quadrant in the graph) show strong overlap, with 76% of the SPECC1L interactors identified in the MYPT1 dataset and 43% of the MYPT1 interactors identified in the SPECC1L dataset. These overlapping factors, listed in the colour-coded table, are involved in the regulation of cell contractility, actin organization, microtubule stability, junction turnover, cytokinesis, adhesion and migration. A Gene Ontology Enrichment Analysis (Ashburner et al 200, NAR2021) is included in Supp Data File 1.

### MYPT1 directly interacts with SPECC1L via their respective C-termini

Using GlobPlot2.3 (http://globplot.embl.de/)[34] to predict regions of disorder and globularity, we designed a truncation mutant strategy to probe for the association of specific regions of both proteins and determine whether or not their interaction is direct. We first assessed co-precipitation of endogenous MYPT1 from cell lysates with GFP-tagged SPECC1L truncation mutants (Fig. 2a). Our results indicated that the C-terminal half of SPECC1L (SPECC1L-CT; aa 462-1117) mediates its association with MYPT1, as confirmed both by IP/WB (Figure 2b) and BioID (Fig. S1c). Similarly, the C-terminal half (aa 462-1068) of the related family member SPECC1 governs its association with MYPT1 (Figure S1f). Further truncation of the C-terminus of SPECC1L (ΔNTΔCHD) revealed that the C-terminal actin-binding calponin homology (CH) domain is not required for MYPT1 association (Fig. 2b).

**Figure 2.**
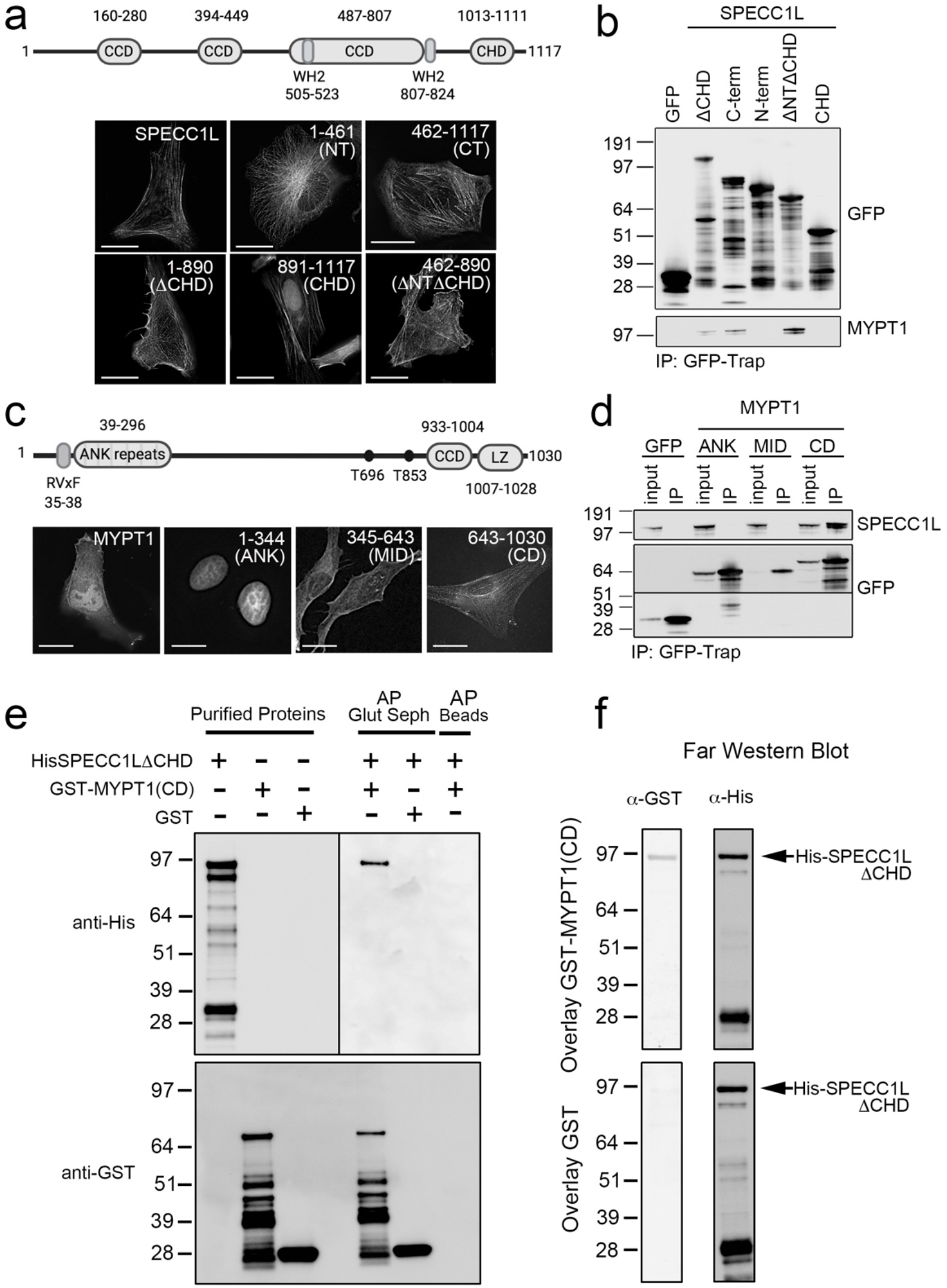
SPECC1L binds MYPT1 directly. a. SPECC1L contains 3 coiled-coil domains (CCD), a C-terminal calponin homology domain (CHD) and 2 putative WASP homology domain-2 (WH2) actin binding domains. Various truncation mutants exhibit distinct subcellular localizations. b. The minimal region of SPECC1L required for association with MYPT1 is ΔNTΔCHD (aa 462-890), which contains the third CCD. c. MYPT1 consists of a series of N-terminal ankyrin repeats adjacent to an RVxF PP1β binding motif and C-terminal coiled coil and leucine zipper (LZ) domains. Three truncation mutants (ANK, aa 1-344; MID, aa 345-643; CD, aa 643-1030) based on the literature demonstrate the expected subcellular localization patterns. d. The minimal region of MYPT1 required for association with SPECC1L is the CD region, which includes the CCD and LZ domain. (e-f) Recombinant His-SPECC1LΔNTΔCHD binds recombinant GST-MYPT1(CD) directly in both in vitro co-precipitation (e) and far Western blot (f) assays. GST alone is used as a negative control for both. Scale bars are 10 μm.

To determine which region(s) of MYPT1 mediate its association with SPECC1L, we divided the protein into three fragments: an N-terminal region containing the PP1-binding RVxF motif followed by a series of ankyrin repeats (ANK; aa 1-344), a middle region (MID; aa 345-653) and a C-terminal region that contains a coiled-coil domain (CCD) and leucine zipper (LZ) domain (CD; aa 654-1030) (Fig. 2c). The localization of the fragments is consistent with previous reports that show GFP-MYPT(ANK) to be nuclear, (MID)Mypt1-GFP to be both nuclear and cytoplasmic and GFP-Mypt1(CD) to be predominantly cytoplasmic in association with the cytoskeleton [35]. When affinity purified from cell lysates, only MYPT1(CD) co-purified endogenous SPECC1L (Fig. 2d). This region also mediates association with known interactors that include myosin, the active form of RhoA, the M20 myosin phosphatase subunit and the inhibitory CPI-17 protein (see [22] for review). Finally, we confirmed that, when co-expressed in cells, GFP-MYPT1(CD) co-purifies mCherry-tagged SPECC1L-CT (Fig. S1d).

We next set out to determine if SPECC1L directly binds MYPT1. To do this, we first expressed and purified recombinant GST-MYPT1(CD). Recombinant SPECC1L-CT was more susceptible to degradation, so we tested all of the fragments and chose to work with SPECC1LΔCHD (1-890), which is a close representation of the full-length protein that retains MYPT1 association (Fig. 2b). Recombinant His-SPECC1LΔCHD was mixed with recombinant GST-MYPT1(CD) or GST alone, which were then captured on Glutathione Agarose beads for detection of co-purified HisSpecc1LΔCHD by WB analysis using anti-His antibodies (Fig. 2e). This in vitro co-precipitation assay confirmed direct and specific interaction of His-SPECC1LLΔCHD with GST-MYPT1(CD). In a complementary far Western blot approach, purified His-SPECC1LΔCHD was resolved on a 1D SDS-PAGE gel, transferred to a nitrocellulose membrane and overlaid with either purified GST-MYPT1(CD) or GST alone. Fig. 2f shows the specific and direct binding of GST-MYPT1(CD) to His-SPECC1LΔCHD.

### Distinct regions of SPECC1L mediate its association with microtubules and actin filaments

The distinct subcellular localizations observed for the various SPECC1L truncation mutants (Fig. 2a) were consistent with previous results suggesting SPECC1L associates with both the microtubule (MT) network and the actin cytoskeleton [25]. We first confirmed that the network at which the N-terminal half (SPECC1L-NT; aa 1-461) accumulates counter-stained with anti-α-tubulin (Fig. 3a). This localization pattern was lost when cells were treated with the polymerization inhibitor nocodazole (NOC) to disrupt the MT network (Fig. 3a, bottom panels). We therefore concluded that this region of SPECC1L mediates MT association. Similarly, the N-terminal half of SPECC1 also governs its association with MTs (Fig. S2c).

**Figure 3.**
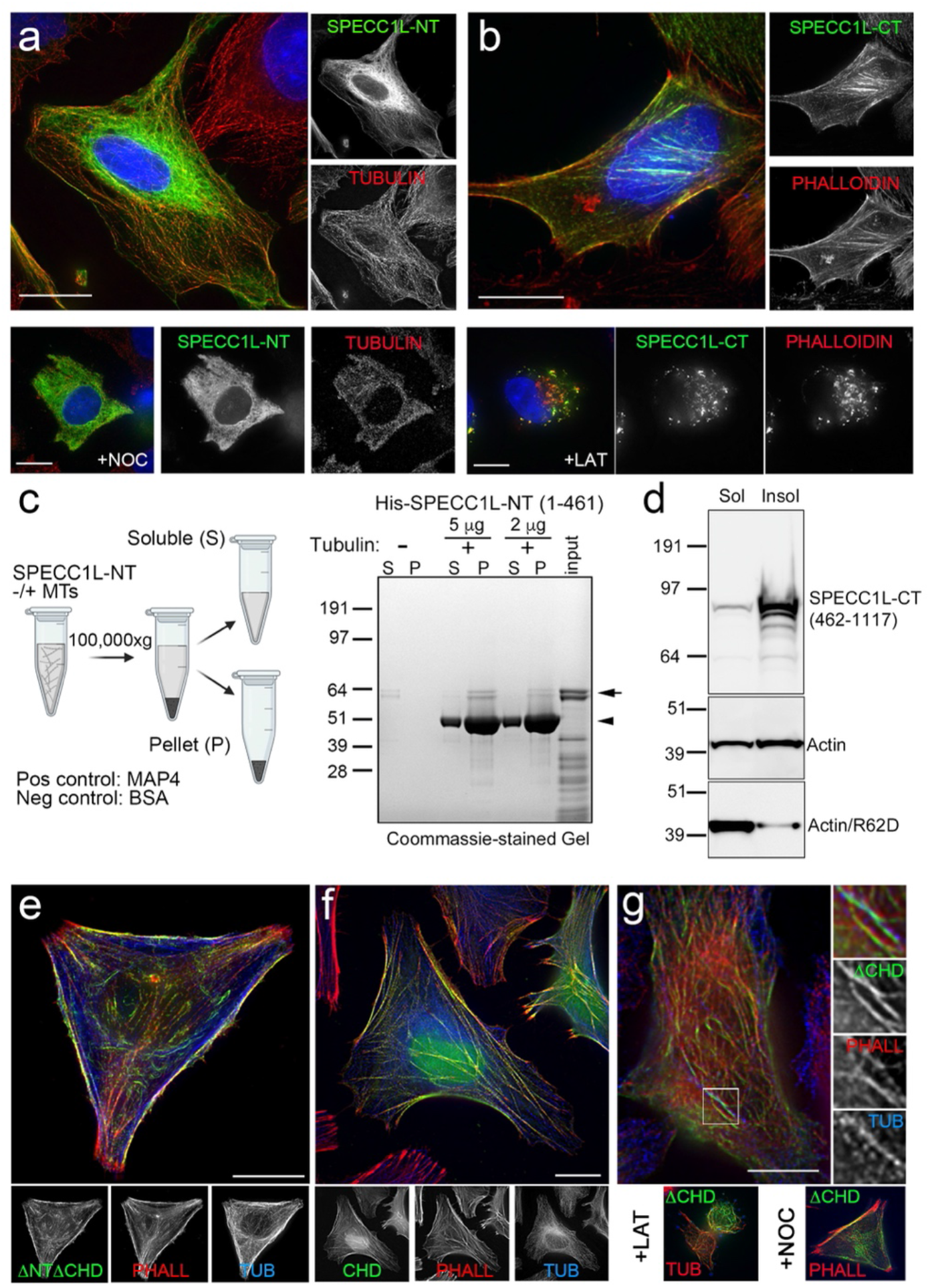
Distinct regions of SPECC1L mediate its association with microtubules and actin filaments. a. The N-terminal half (NT; aa 1-461; green) of SPECC1L associates with microtubules that counterstain with anti-alpha-tubulin-AlexaFluor 568 (red) in U2OS cells, and this localization is disrupted with nocodazole (NOC) treatment (8.3 uM for 2 hrs). Hoechst 33342-stained DNA is shown in blue. B. The C-terminal half (CT; aa 461-1117; green) of SPECC1L associates with actin filaments that counterstain with AlexaFluor568-phalloidin (red), and this localization is disrupted with latrunculin B (LAT) treatment (2 uM for 2 hrs). Hoechst 33342-stained DNA is shown in blue. c. Purified SPECC1L-NT binds microtubules directly in a microtubule binding assay, showing an enrichment (arrow) in the pellet (P) vs. the supernatant (S) fraction when tubulin (arrowhead) is included in the assay. Controls and WB detection of His-SPECC1L and Tubulin are in Fig. S2. d. In an actin fractionation assay, the majority of SPECC1L-CT remains in the insoluble fraction with filamentous actin when U2OS cells are extracted with Triton X-100. The nonpolymerizable Actin R62D mutant, which remains soluble, is included as a negative control. SPECC1L was detected using anti-GFP antibodies and Flag-Actin (WT and R62D) was detected using anti-Flag antibodies. SPECC1L ΔNTΔCHD (e) and SPECC1L-CHD (f) both co-localize with phalloidin-stained F-actin structures (red) in U2OS cells, albeit with different localization patterns. Tubulin staining is shown in blue. g. SPECC1L lacking the CHD (green) shows overlap with both stained microtubules (anti-tubulin; blue) and actin filaments (phalloidin; red), and demonstrates both a LAT-resistant MT pool and a NOC-resistant actin filament pool. Scale bars are 10 μm.

In order to test whether the association of SPECC1L with MTs is direct or indirect, we performed a microtubule binding protein spin-down assay. Purified recombinant His-tagged SPECC1L-NT was mixed with freshly prepared MTs and the reaction mix centrifuged at 100,000 x g. At this speed, the MTs pellet along with any protein that directly associates with them. SPECC1L-NT was observed to specifically pellet in the presence of MTs, and remained in the supernatant in their absence (Fig. 3c; Fig. S2b). BSA was included as a negative control, as it does not bind MTs and thus remains in the soluble fraction, while the known MT binding protein MAP4 was included as a positive control (Fig. S2a). This assay confirms, for the first time, that SPECC1L is a bona fide MT binding protein.

The presence of a calponin homology (CH) domain [31] at the C-terminus of SPECC1L (Fig. 2a) has been suggested to facilitate actin binding. Our analysis using the ELM (eukaryotic linear motif) online resource [36] also identified two putative WH2 actin-binding domains in the C-terminal half of SPECC1L, upstream of the CH domain (Fig. 2a). We first confirmed that the structures at which SPECC1L-CT accumulates counterstain with fluorophore-tagged phalloidin, a high-affinity probe for F-actin (Fig. 3b). This localization pattern was lost when cells were treated with Latrunculin B (LAT), which sequesters monomeric G-actin and induces disassembly of actin filaments. Using a classic actin fractionation approach [37], we further demonstrated that GFP-tagged SPECC1L-CT expressed in U2OS cells is resistant to Triton X-100 extraction, remaining in the insoluble cytoskeletal fraction with F-actin (Fig. 3d). The nonpolymerizable actin R62D mutant[38] was included to demonstrate its shift to soluble (G-actin) pools.

To test the contributions of the predicted actin binding regions to this localization, we removed either the CH domain (ΔNTΔCHD; aa 461-890) or the WH2 domains (CHD; aa 890-1117) from the C-terminal half of SPECC1L. Removal of the CH domain did not obviate actin association (Fig. 3e), although the pattern differs from that observed for the CH domain-containing fragment (Fig. 3f). While the CHD derivative distributes along straight stretches of a filamentous network, the ΔNTΔCHD mutant associates with both cortical filaments and in shorter structures in the cytoplasm. Both localization patterns are disrupted with LAT but not NOC treatment (Fig. S2c-d), confirming that they represent accumulations at actin structures. This suggests that both regions play roles in the targeting of SPECC1L to the actin network, possibly by mediating association with specific actin structures.

### Overexpression of SPECC1L induces stabilized and acetylated microtubule bundles

Consistent with previous reports in the literature [25][39], immunofluorescence analysis of endogenous SPECC1L shows overlap with the MT and F-actin networks during interphase, although the latter predominates (Fig. 4a). This was also observed for SPECC1L-GFP stably overexpressed at endogenous levels in U2OS cells (Fig. 4b). Interestingly, removing the CH domain shifts this pattern, with the SPECC1L-ΔCHD truncation mutant demonstrating both filamentous-actin and MT pools (Fig. 3g). This suggests a dynamic balance of distribution between these two networks that is regulated, at least in part, by competition between the MT binding region and the actin association domains.

**Figure 4.**
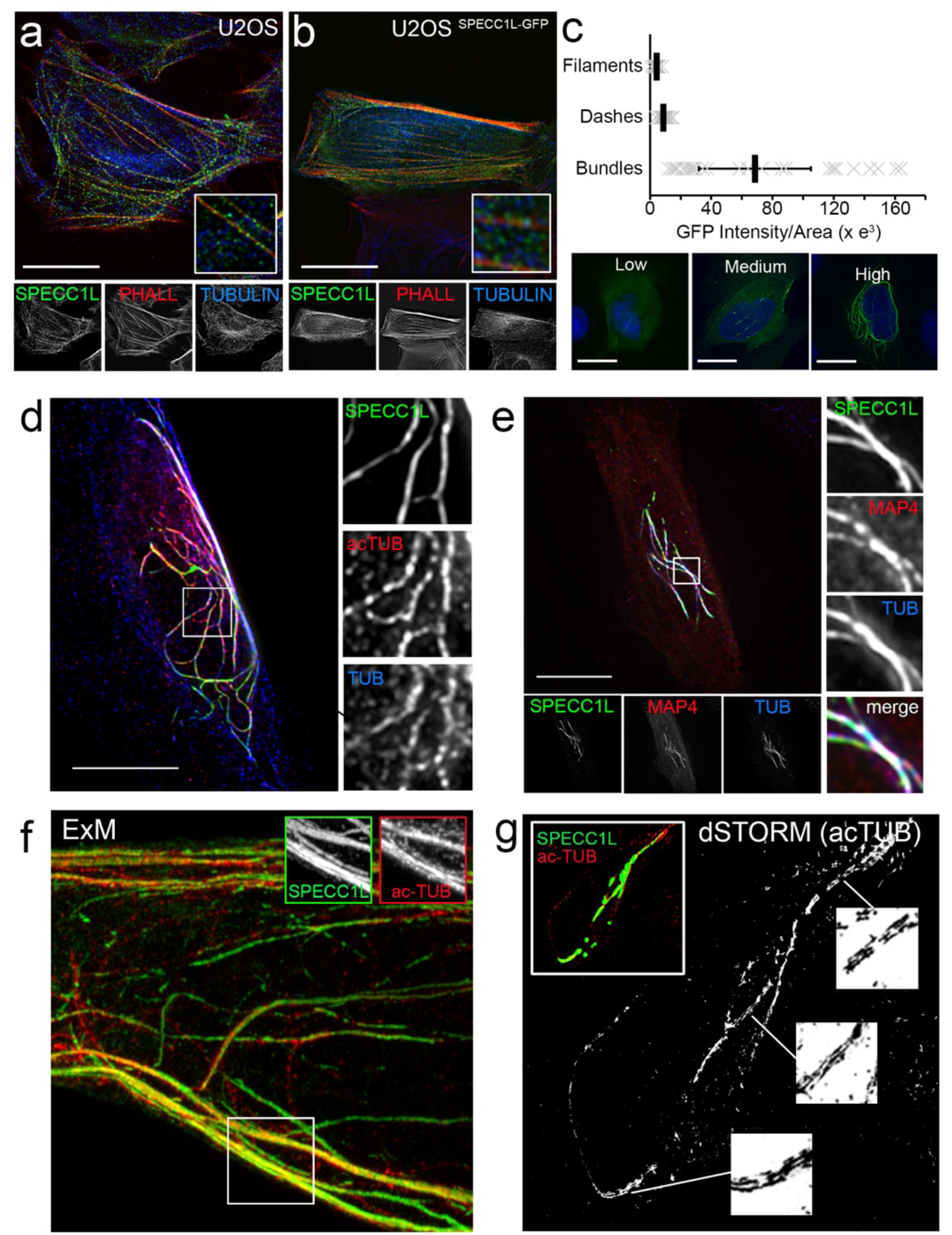
Overexpression of SPECC1L induces stabilized and acetylated microtubule bundles. a. In untreated interphase U2OS cells, the localization of endogenous SPECC1L detected by immunostaining (green) is predominantly cytoplasmic and overlaps the pshallodin-stained F-actin pattern (red) more closely than the MT pattern detected by anti-alpha-tubulin staining (blue). b. A similar pattern is observed for SPECC1L-GFP stably expressed at endogenous levels in the U2OS^SPECC1L-GFP^ cell line. c. Overexpression of SPECC1L at increasingly higher levels shifts localization from predominantly filamentous to accumulation in cytoplasmic “dashes” (at medium intensities) and “bundles” (at high intensities). GFP intensities and localization were annotated for >60 cells in 3 independent experiments (bar = mean ± SE). d. Cytoplasmic SPECC1L bundles (green) counterstain with antibodies against both alpha-tubulin (blue) and alpha-tubulin acetylated on Lys40 (acTUB; red). e. mCh-MAP4 (red) accumulates at SPECC1L-GFP (green) MT bundles counterstained with anti-alpha-tubulin (blue). Analysis of these SPECC1L/acetylated MT bundles at nanoscale resolution by expansion microscopy (ExM; f) and dSTORM single molecule localization microscopy (g) confirmed that the thick structures consist of multiple MTs aligned in parallel and in close association. Scale bars are 10 μm.

Expression of increasingly higher levels of full-length SPECC1L resulted in aberrant accumulation in thick, thread-like structures (Fig. 4c). These structures counterstain with antibodies that detect both alpha-tubulin or its acetylated (on K40) form, the latter of which is a marker for stabilized MTs (Fig. 4d; [40]). They are also sites of accumulation of the Tau family Microtubule Associated Protein 4 (MAP4; Fig. 4e), which has been shown to promote MT stabilization and regulate MT-based motility [41].

The thickness of the structures suggested bundling of multiple MTs, which we assessed by super-resolution imaging. First, we used a 10X Expansion Microscopy approach to isotropically expand U2OS cells transiently overexpressing SPECC1L-GFP and fixed and stained with anti-GFP (to boost the SPECC1L signal) and anti-acetylated-alpha-tubulin antibodies [42], [43]). Airyscan-based imaging of gel segments using a Zeiss LSM880 confocal laser scanning microscope confirmed that the thick structures represent multiple closely-associated MT strands lying in parallel (Fig. 4f). This was also observed when we used the complementary dSTORM (direct Stochastic Optical Reconstruction Microscopy) super-resolution imaging approach to visualize the acetylated tubulin signal in regions of SPECC1L-GFP accumulation (Fig. 4g). Interestingly, we did occasionally observe transient accumulation of SPECC1L-GFP at MT bundles in our stable cell line, with live imaging showing that it could reverse over time (Fig. S3c). This is consistent with the idea that a localized elevated level of SPECC1L, which can associate with both the actin cytoskeleton and the microtubule network, has the potential to impact their homeostatic balance in cells.

It is important to note that this MT bundling phenotype is only observed with overexpression of full-length SPECC1L. Even at very high levels of over-expression, the MT binding region alone (SPECC1L-NT) only accumulates at MTs and does not induce bundling, suggesting contributions from the actin and/or MYPT1 association domains to this phenotype. Consistent with this, although removal of the C-terminal actin binding CH domain (SPECC1L-ΔCHD) promotes increased association with MTs (Fig. 3g), the thick, bundled MT phenotype is not observed with overexpression of this mutant [25].

Sequence alignment of the N-terminal regions (aa 1-461) in SPECC1L and SPECC1 identified a stretch within the second coiled coil domain that shows a high degree of similarity (Fig. S5c). Mutations in this region in SPECC1L have been identified in patients with the congenital developmental disorders Opitz G/BBB and Teebi hypertelorism [26]–[29] (Fig. 7a), and previous studies suggested that disrupted MT association may contribute to the observed phenotypes. Although we did not detect changes in MT association or the stabilization/bundling phenotype for two of these mutations (T397P and Q415P) when introduced into SPECC1L or SPECC1L-NT, it may be that the impact is subtle and effects accumulate over time. A more recent study reported diminished overlap with MTs when the entire CCD2 was removed [39]. Functional MT binding may also require additional regions in the N-terminal half of the protein. SPECC1 is expressed as multiple splice variants, and we have compared SPECC1 isoform 1 (NSP5b3b) to SPECC1L as they are the most similar in size and structure. There is, however, a splice variant (isoform 4; NSP5a3b) in which the N-terminal residues 1-94 are replaced by a unique stretch of 13 aa. Interestingly, although SPECC1/iso1 shows the same MT bundling phenotype as SPECC1L (Fig. S5a), SPECC1/iso4 remains predominantly associated with actin filaments, even at high levels of overexpression (Fig. S5b). Although SPECC1 contains a potential KXGS MT binding domain within its unique N-terminus, similar to those found in the MT binding domains of Tau, MAP2C and MAP4 (Figure S5c), fusion of the 94 aa region to GFP did not promote MT association.

### SPECC1L modulates the subcellular distribution and turnover dynamics of the myosin phosphatase complex

To determine whether overexpression of full-length SPECC1L can also shift the balance MYPT1 localization, we stained cells transiently overexpressing SPECC1L-GFP with anti-MYPT1 antibodies and confirmed that a pool of endogenous MYPT1 is recruited to the SPECC1L-containing MT bundles (Fig. S4a). We next assessed effects on the localization and turnover dynamics of MYPT1 in a HeLa Bacterial Artificial Chromosome (BAC) cell line that drives expression of GFP-tagged mouse Mypt1 (91.7% identical to the human protein) under the control of its endogenous promoter [44]. For these experiments, we expressed BirA*-tagged SPECC1L, which allowed us to compare the localization of Mypt1-GFP to the proximity labeling profile of SPECC1L detected with fluorophore-tagged streptavidin. This confirmed that all cells that showed a bundled MT phenotype withr Mypt-GFP were also expressing the SPECC1L fusion protein (Fig. 5a, top panel). Similarly, co-expression of SPECC1L-CT induced a visible increase in accumulation of Mypt1-GFP on F-actin (Fig. 5a, bottom panel).

**Figure 5.**
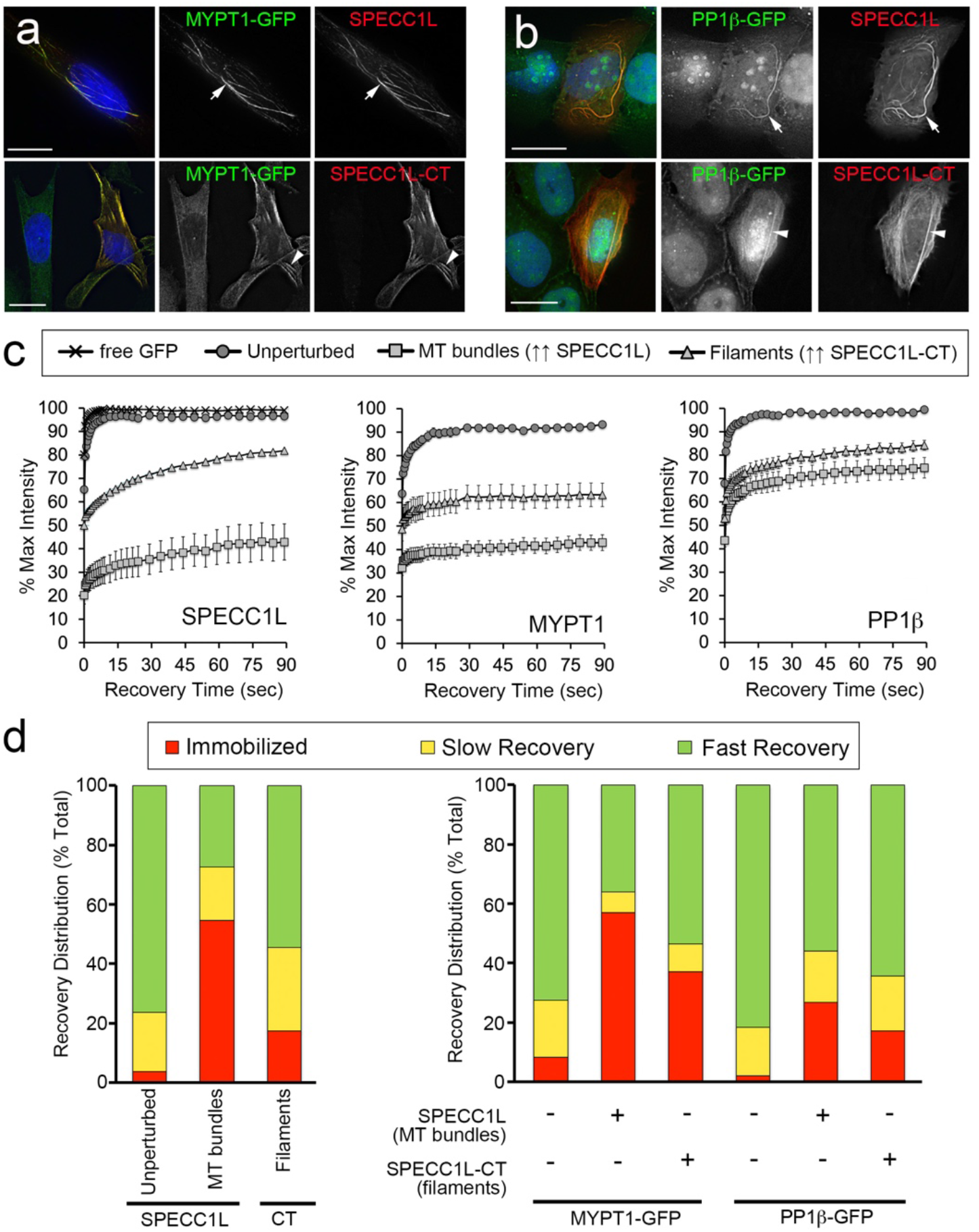
SPECC1L modulates the subcellular distribution and turnover dynamics of the myosin phosphatase complex. a. Mypt1-GFP (green) in the BAC HeLa line is recruited to MT bundles induced by transient overexpression of SPECC1L-BirA* (top panel, arrows), and to actin filaments at which transiently overexpressed SPECC1L-CT accumulates (lower panel, arrowheads). Biotinylation patterns were detected using AlexaFluor 647-tagged Streptavidin (red). Hoechst 33342-stained DNA is shown in blue. b. PP1β-GFP (green) in the U2OS^PP1β-GFP^ stable line is recruited to MT bundles induced by transient overexpression of SPECC1L (top panel, arrows) and to actin stress fibers at which transiently overexpressed SPECC1L-CT accumulates (lower panel, arrowheads). Biotinylation patterns were detected using AlexaFluor 647-tagged Streptavidin (red). Hoechst 33342-stained DNA is shown in blue. c. Fluorescence Recovery After Photobleaching (FRAP) curves for GFP-tagged SPECC1L- and SPECC1L-CT (unperturbed represents SPECC1L-GFP in the U2OS ^SPECC1L-GFP^ stable line) and for Mypt1-GFP and PP1β-GFP either unperturbed or at the structures to which they are recruited by overexpressed SPECC1L or SPECC1L-CT (as shown in panels a-b). d. Tables summarizing the FRAP results, based on mobile fractions and recovery times calculated using GraphPad Prism (Table S1). In all cases, the recovery curves for the mobile fraction were best fit by a double exponential line, indicating two pools with different recovery times (slow and fast). Representative FRAP experiments are shown in Fig. S4. Scale bars are 10 μm.

Given that SPECC1L co-purifies MYPT1/PP1β as a complex (Fig. 1d; Supp Data File 1), we reasoned that SPECC1L-induced relocalization of Mypt1 would be accompanied by a concomitant change in the localization of PP1β. For these experiments we utilized a U2OS cell line that stably expresses a low level of GFP-tagged PP1β. We demonstrated that overexpression of full-length SPECC1L recruits a pool of PP1β-GFP to bundled MTs (Fig. 5b, top panel), while overexpression of the C-terminal half recruits excess PP1β-GFP to actin filaments (Fig. 5b, bottom panel).

Having confirmed that relocalization of both GFP-tagged Mypt1 and PP1β is sufficiently dramatic to allow us to unambiguously identify non fluorophore-tagged SPECC1L and SPECC1L-CT expressing cells based on their altered subcellular localization patterns, we set out to assess their turnover dynamics by Fluorescence Recovery After Photobleaching (FRAP) analysis in live cells. In these experiments, the pool of GFP in a region of interest (ROI) in the cell is irreversibly photobleached using a high-powered 488 nm laser, and the return of a GFP signal to this ROI (which represents exchange with pools of GFP outside the ROI) monitored over time (Fig. S4b). Recovery profiles that are slower and less complete than that observed for free GFP over the same time scale suggest association with underlying subcellular structures/complexes.

Analysis of cytoplasmic SPECC1L-GFP in the stable cell line revealed that the fusion protein is relatively dynamic, albeit with slower turnover dynamics compared to GFP alone (Fig. 5c, left graph, circles vs crosses). Overexpressed SPECC1L-GFP that accumulated at bundled MTs has a significantly reduced mobile fraction with a slower recovery rate (Fig. 5c, left graph, squares), suggesting near immobilization at these structures. As previously noted, the N-terminal half of SPECC1L, when overexpressed as a GFP fusion in cells, accumulates on the MT network (Fig. 3a) but does not induce bundling/stabilization. Assessment of its turnover dynamics by FRAP revealed full recovery within 90 sec post-bleach, and a slower recovery rate than free GFP, suggesting a normal on/off association with MTs (not shown). This again suggests that the bundling/stabilization phenotype requires a specific region within the C-terminal half of the protein.

Mypt1-GFP also showed a reduced mobile fraction and slower turnover rate compared to free GFP, confirming the presence of underlying binding events (Fig. 5c, middle graph, circles). Overexpression of full-length SPECC1L induced accumulation of a pool of Mypt1-GFP at bundled MTs and a dramatic reduction in its turnover dynamics at these structures (Fig. 5c, middle graph, squares), similar to the near immobilization observed for SPECC1L-GFP. PP1β-GFP also showed slower turnover dynamics than free GFP (Fig. 5c, right graph, circles), and its mobility changed significantly when it was recruited to MT bundles by overexpressed full-length SPECC1L (Fig. 5c, right graph, squares). Representative FRAP experiments for SPECC1L-GFP, Mypt1-GFP and PP1β-GFP at MT bundles are show in Fig. S4c.

FRAP analysis of the C-terminal SPECC1L fragment, which accumulates at actin filaments (Fig. 3), also showed a significantly reduced mobile fraction and slower turnover rate (Fig. 5c, left graph, triangles). Consistent with this observation, reduced mobile fractions and slower turnover rates were observed for the pools of excess Mypt1-GFP (Fig. 5c, middle graph, triangles), and PP1β-GFP (Fig. 5c, right graph, triangles) that were recruited to actin filaments by overexpression of SPECC1-CT. Representative FRAP experiments for SPECC1L-GFP, Mypt1-GFP and PP1β-GFP at actin filaments are show in Fig. S4d. It should be noted that PP1β did not show the same degree of reduced mobility as MYPT1 at the MT bundles and actin filaments. This is not unexpected, however, as it may be that SPECC1L and MYPT1 are more tightly associated with these structures while PP1β can still turn over on MYPT1 as it exchanges with other PP1 holoenzyme complexes.

While the recovery curves allow direct visual comparison by demonstrating the changes in turnover dynamics qualitatively, we wanted to quantitatively assess these changes by calculating mobile fractions and recovery half-times. We were unable to fit single exponential recovery curves to the data, indicating the presence of more than one recovery pool. Double exponential curves provided a better fit in each case (R_2_ values > 0.95). For all conditions, we determined the fraction that does not recover within 90 sec post-bleach (immobilized), the fraction that recovers with fast kinetics and the fraction that recovers with slow kinetics. The half-times of recovery for the fast and slow pools were also determined. These values are summarized in Table 1, with the distribution of the GFP fusion proteins between immobile/slow/fast pools graphically represented in Fig. 5d. Taken together with the observed changes in subcellular distribution (Fig. 5a-b), they show that SPECC1L has a profound effect on both the localization and turnover dynamics of MYPT1 and PP1β. These results suggest that SPECC1L can directly mediate the homeostatic distribution of MYPT1/PP1β between the MT and F-actin networks in interphase cells (Fig. 7b).

### Association of SPECC1L with actin- and microtubule-related proteins is dynamic and changes throughout the cell cycle

Our MYPT1 and SPECC1L interactomes detected several actin binding proteins that also associate with CTs and can regulated their stability. These include CAPZB and Coronin and Spectrin family members (Fig. 1e; Supp Data File 1). MT associated proteins (MAP1A/B, MAP4) were also detected but not as highly enriched (Supp Data File 1). When we used a BioID/MS approach to map the interactome of transiently overexpressed SPECC1L-BirA*, we saw a significant enrichment of MAP4 (Fig. 6a) that is consistent with its observed accumulation at SPECC1L-induced MT bundles (Fig. 4e). The MYPT1/PP1β complex was also highly enriched in the BioID dataset, along with several structure and motility-related factors that include myosin (MYO1B), cortactin (CTTN), Coronin-1B (COROB1B) and the CRK-CRKL adaptor proteins that bridge tyrosine phosphorylation events to diverse intracellular signalling pathways (Supp Data File 2).

**Figure 6.**
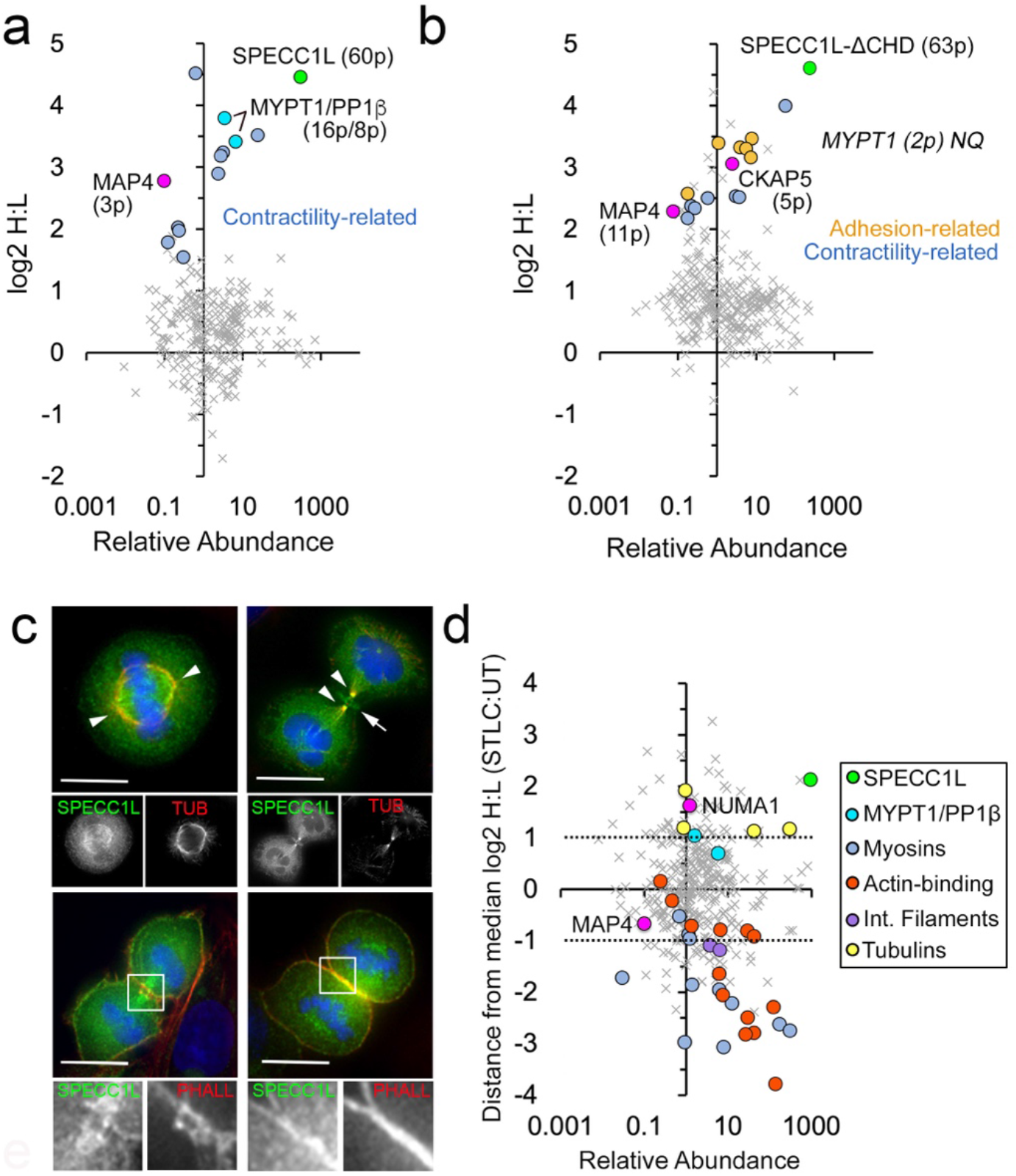
Association of SPECC1L with both microtubule and actin cytoskeletal proteins is dynamic and changes throughout the cell cycle. a. The graph shows proteins that are biotinylated by BirA*-tagged SPECC1L in a quantitative SILAC BioID/MS experiment. For the highlighted proteins, the number of peptides detected is indicated in parentheses. b. The graph shows proteins that are biotinylated and enriched by BirA*-tagged SPECC1L-ΔCHD in a quantitative SILAC BioID/MS experiment. For the highlighted proteins, the number of peptides detected is indicated in parentheses. As noted, MYPT1 was detected but not quantified (summed heavy:light peptide intensities were 47:1). The full BioID/MS datasets are provided in Supp Data File 2. c. The top panels show co-localization of SPECC1L (green) in the U2OS^SPECC1L-GFP^ stable line with spindles (left, arrowheads in metaphase cell) and midbodies (right, arrowheads in telophase cell) visualized by anti-tubulin staining (red). Hoechst 33342-stained DNA is shown in blue. Additional accumulation between the midbodies was observed in telophase cells (arrow). The bottom panels show co-localization of SPECC1L (green) in the U2OS^SPECC1L-GFP^ stable line at the cleavage furrow and the cortex with Phalloidin-stained actin filaments (red). Hoechst 33342-stained DNA is shown in blue. d. Results of a quantitative SILAC AP/MS experiment comparing enrichment of interactors with SPECC1L-GFP immunoprecipitated from asynchronous U2OS^SPECC1L-GFP^ cells (L) vs. U2OS^SPECC1L-GFP^ cells arrested at metaphase by overnight treatment with S-trityl-L-cysteine (STLC; H). Enrichment is expressed as the displacement from the median H:L ratio, with factors enriched < 2-fold in either direction considered to bind equally under both conditions (area between the dashed lines). Interactors enriched >2-fold with SPECC1L-GFP from mitotic cell extracts are above the dashed line while interactors enriched >2-fold with SPECC1L-GFP from asynchronous cell extracts fall below the dashed line. The full AP/MS dataset is provided in Supplemental Data File 1. Scale bars are 10 μm.

Having observed that removal of the C-terminal CH domain shifts the balance of steady-state SPECC1L association more equally between MTs and actin filaments (Fig. 3g), we were interested to see how that was reflected in its proximity interactome. As shown in Fig. 6b, MAP4 is again enriched, along with the MT polymerase CKAP5[45](Fig. 6b). The Coronins CORO1B and CORO1C were also enriched, as were adhesion-related Syndecan family proteins (SDC1/2/4) that provide mechanical links between the extracellular matrix and the actin cytoskeleton.

Another example of a shift of SPECC1L from actin to tubulin structures is the mitotic spindle association observed in early mitosis (Fig. 6c). We reasoned that this would be reflected in its interactome, and quantitatively compared proteins enriched with SPECC1L-GFP (by SILAC AP/MS) from asynchronous U2OS^SPECC1L-GFP^ cells (primarily interphase) vs U2OS^SPECC1L-GFP^ cells arrested at metaphase by overnight treatment with the Eg5 inhibitor S-trityl-L-cysteine (STLC). The graph in Fig. 6d shows the distance from the median log2 H:L ratio for all identified proteins (vs their relative abundance in the AP), highlighting those were enriched >2-fold in either the H (metaphase; above the top dashed line) or L (interphase; below the bottom dashed line) condition. Proteins in the middle region between the 2 dashed lines were equally enriched in both conditions. Association of SPECC1L with MYPT1/PP1β persists throughout the cell cycle, which was further confirmed by AP/WB analysis (Fig. S6b).

Consistent with its change in localization, the majority of actomyosin-related proteins were enriched more with SPECC1L from asynchronous cell lysates, while STLC treatment led to an increased association with tubulin. The centrosomal NuMA (nuclear mitotic apparatus) protein, which interacts with MTs and play a role in the formation and organization of the mitotic spindle, was also enriched with SPECC1L in STLC-arrested cells. Along with dynein, this protein is required to anchor minus-end microtubules to centrosomes in mitosis [46] and promotes spindle bipolarity by organizing the radial array of MTs that incorporates Eg5 [47]. When we imaged SPECC1L-GFP in STLC-arrested cells, we observed a clustering at the center of the monoaster spindle (Fig. S6a) similar to that shown previously for NuMA[47], [48]. This suggests that they associate at the spindle pole. Transient recruitment of SPECC1L to centrosomes had been previously noted but not shown. Long-term live imaging of our U2OS^SPECC1L-GFP^ cell line confirmed enrichment of GFP-tagged Specc1L at centrosomes just prior to the onset of mitosis (Fig. S6c-d). At later stages of mitosis, SPECC1L remains associated with MTs at the midbody and shows additional accumulation at phalloidin-stained actin filaments at the cortex and the cleavage furrow (Fig. 6c).

## DISCUSSION

MP was one of the first Ser/Thr protein phosphatase complexes identified [49], and a wealth of literature over the past three decades has functionally dissected its critical role in the regulation of actomyosin contractility. Along the way, there has been a growing appreciation of its contribution to the regulation of other key cellular events, including cytoskeletal organization, adhesion, mitosis and transcription (see [22] for review). Its interactors are found throughout the cell, but what remains unclear is how it balances these diverse functions (and its subcellular distribution) and adapts to physiological changes. sAs an example, a recent elegant study[24] identified HDAC6, which plays an important role in microtubule deacetylation, as a substrate for MYPT1/PP1β. Our data are consistent with their model, which suggests the MP complex balances contractility and microtubule acetylation by associating either with myosin light chain or HDAC6 (Fig. 7). What has remained to be answered is how MYPT1 is efficiently recruited to these substrates in the cell, and what drives association with one over the other.

**Figure 7.**
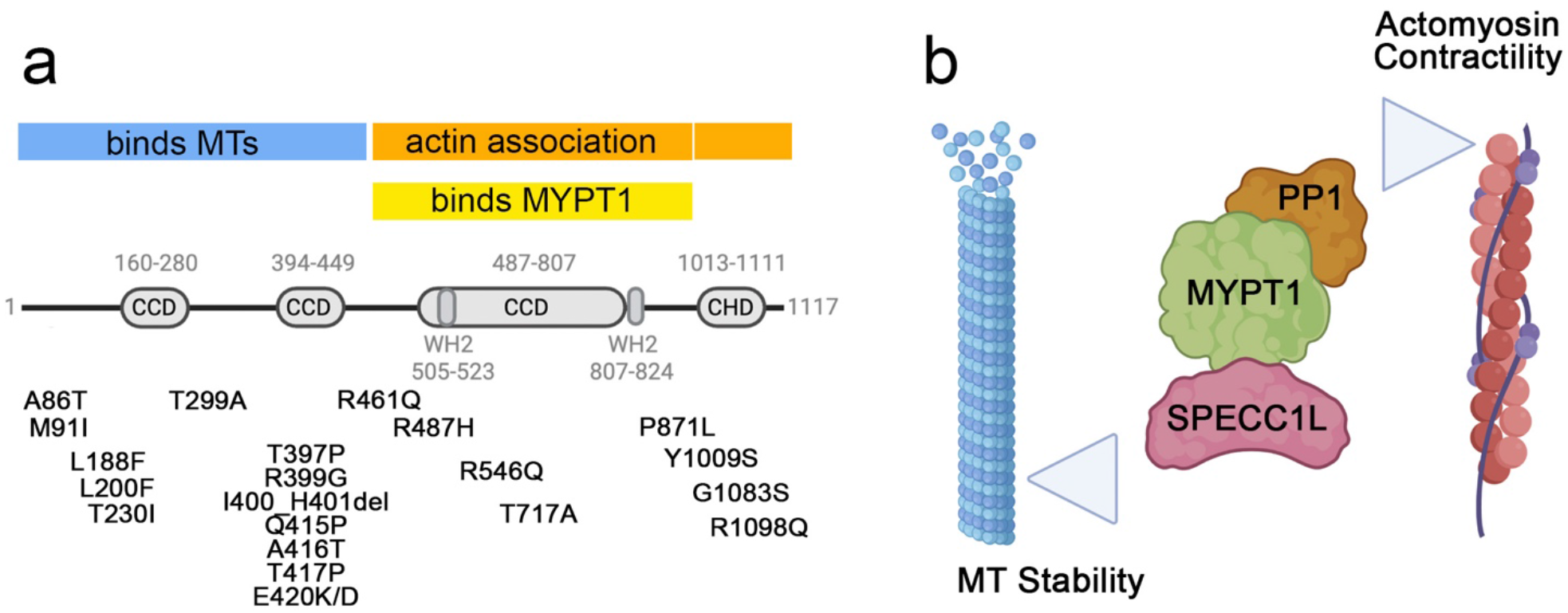
Location of disorder-related mutations within SPECC1L. a. Summary of SPECC1L mutations that have been linked to developmental defects. b. Our proposed model in which SPECC1L regulates the balance of MYPT1/PP1β activity between MT and actomyosin-associated substrates throughout the cell cycle via direct binding to MYPT1.

Our identification of a novel complex comprising Mypt1/PP1β and proteins capable of associating dynamically with both the MT and actin networks makes SPECC1L (and possibly SPECC1) an attractive candidate for the “missing link” that regulates the distribution of phosphatase activity between these two structures. We showed that SPECC1L associates with MTs and with F-actin via N-terminal and C-terminal regions, respectively, and that it directly binds MYPT1 in a stable complex that can be purified from cell extracts. We also showed significant overlap of their interaction profiles under steady-state conditions. Furthermore, we demonstrated that SPECC1L can regulate the subcellular localization and turnover dynamics of the MYPT1/PP1β complex in cells, pushing the balance more toward MTs (and a stabilized/bundled phenotype) or F-actin (and increased stress fibers). Although HDAC6 was not detected in our SPECC1L BioID interactome (and was close to threshold in our SPECC1L AP/MS interactome), that does not rule out their association as it was not detected in our MYPT1 interactome either. The MT protein enriched the most in our SPECC1L BioID experiments (both full-length and the ΔCHD mutant) was MAP4. MAP4 was also detected, albeit closer to threshold, in our MYPT1 and SPECC1L AP/MS datasets. With phosphorylation inducing its detachment from MTs, which in turn are destabilized [50], our future work will explore the possibility that MAP4 is a substrate for MYPT1/PP1β targeted to MTs by excess SPECC1L.

An interesting observation was that the single CH domain of SPECC1L can mediate, at least in part, its association with actin, as two tandem CH domains are normally required for this interaction. A previous study noted that the CH domain in SPECC1L showed moderate homology to those found in α-actinin, Macrophin, Beta-Spectrin and Smoothelin [30]. Interestingly, Smoothelin-like protein SMTNL1, which is primarily expressed in smooth and striated muscle cells, has been shown to be a negative regulator of both MP activity (toward myosin light chain) and MYPT1 expression [51][52]. We saw no indication that overexpression of SPECC1L altered MYPT1 protein levels, but future work will address whether or not it can modulate MP activity. Given that knockdown of MYPT1 has been shown to induce stress fibers [53], it may be that the stress fibers induced by SPECC1L-CT overexpression contain inactive MYPT1/PP1β. Another obvious question is whether or not SPECC1L itself is a substrate for MP. Although we mapped a residue in SPECC1L (S384) that showed decreased phosphorylation when MYPT1 was overexpressed, neither S384A (non-phosphorylatable) nor S384D (phosphomimic) mutations showed an obvious impact on MT localization or the stabilization/bundling phenotype. More work will be needed to dissect both the functional relevance of the phosphorylation and whether or not it is directly regulated by MP.

Identification of NuMA in the SPECC1L mitotic interactome was consistent with the observed accumulation of SPECC1L at the spindle pole in STLC treated cells. NuMA is a key factor in the formation and organization of the mitotic spindle [54]. It is a structural hub that associates with both microtubules and dynein (among other factors), and binding to dynein has been shown to drive spindle pole focusing and positioning. The dynein/dynactin plus end-directed motor protein complex has also been implicated in mediation of mitotic checkpoint silencing, via transportation of kinetochore checkpoint proteins toward the spindle poles [55]. Interestingly, several dynactins were detected in our MYPT1, but not SPECC1L datasets. This could indicate either that their association represents a MYPT1 role that does not involve SPECC1L, or that MYPT1 forms a tighter complex with them. NuMA was also enriched in our MYPT1 interactome, and both proteins were recently shown to be regulated at centrosomes by O-linked N-acetylglucosamine (O-GlcNAc) modification [56]. It is tempting to speculate that SPECC1L, which we have shown to localize to centrosomes just prior to the onset of mitosis and accumulate at the pole following STLC arrest, also plays a role in the MYPT1-based regulation of spindle formation/maintenance. The specific enrichment of beta-tubulin with SPECC1L in STLC-arrested cells is also intriguing, as its phosphorylation by CDK1 impairs association with MTs [57]. And, given the presence of both MYPT1 and SPECC1L at the cleavage furrow during cytokinesis, future work with synchronized cells will be needed to more comprehensively map their interactomes throughout the cell cycle. Importantly, our work has identified a previously unknown direct interaction of SPECC1L/1 with MYPT1, which can impact the subcellular localization and activity of the MYPT1/PP1β complex. Further characterization of the functional significance of this regulation will be necessary to evaluate its potential as a therapeutic target.

## MATERIALS & METHODS

### Plasmids and antibodies

Full-length MYPT1 and MYPT1 fragments were amplified from cDNA (Open BioSystems) using specific primers and inserted into the pEGFP/mCherry(C1), pEGFP(N3), and pGEX-4-T3 vectors by restriction cloning. Full-length SPECC1L, SPECC1/iso1 and SPECC1/iso4 (and their respective fragments) were inserted into pEGFP(C1), pEGFP(N3), pET-47b(+) and myc-BirA*(C1) and HA-BirA*(N3) vectors using the same approach. All cloning was confirmed by DNA sequencing (StemCore Laboratories, Ottawa Hospital Research Institute). The pEGFP(N3)-PP1β plasmid was previously described [58] and is available through Addgene (plasmid #44223).

Anti-MYPT1 antibodies were obtained from Bethyl Laboratories, anti-SPECC1L(CYTSA) antibodies from ProteinTech and anti-alpha-tubulin, anti-acetylated(K40)-alpha-tubulin and anti-GFP antibodies from Millipore Sigma. The PP1β antibody was previously described [59]. All HRP- and fluorophore-conjugated secondary antibodies and streptavidin were from ThermoFisher.

### Cell culture

U2OS cells were obtained from ATCC and grown in Dulbecco’s modified Eagles’ medium (DMEM) supplemented with 10% fetal calf serum and 100 U/mL penicillin and streptomycin (Wisent Bioproducts Inc). Stable cell lines were generated as previously described and maintained in media supplemented with G418 [60], The mouse Mypt1-GFP Bacterial Artificial Chromosome (BAC) HeLa cell line was a gift from Dr. Tony Hyman (MPI-CBG, Germany)[44]. Cells were transfected with either 1 mg/ml polyethylenimine (PEI; Polysciences, Inc.) or Effectene transfection reagent (Qiagen). For mitotic arrest experiments, cells were treated with either 3.3 uM nocodazole (NOC), or 5 uM S-trityl-L-cysteine (STLC) for 18 hrs. For short-term MT destabilization and disruption of actin polymerization, cells were treated with either 8.3 uM nocodazole (NOC) or 2 uM Latrunculin (LAT) for 2 hrs, respectively.

### Metabolic labeling

Stable isotope labeling with amino acids in cell culture (SILAC) for label-based quantitative MS was carried out as previously described [32]. Briefly, cells were grown for 7-10 passages in high glucose DMEM containing either L-arginine and L-lysine (Light), or the isotopes L-arginine13C/15N and L-lysine13C/15N (Heavy). SILAC media was prepared by supplementing high glucose DMEM minus Arg/Lys/Leu/Met (AthenaES) with 10% dialyzed FBS (ThermoFisher) and the appropriate amino acids, mixing well and filtering through a 0.22 um filter (Millipore). For the SPECC1L metaphase arrest vs asynchronous AP/MS experiment, Light isotope-encoded U2OS^SPECC1L-GFP^ were left untreated for standard harvesting while Heavy-isotope encoded U2OS^SPECC1L-GFP^ cells were treated for 18 hrs with STLC and harvested by mitotic shake-off. For the BioID experiments, Heavy isotope-encoded cells were transfected with BirA*-SPECC1L constructs and Light isotope-encoded cells with the respective GFP-tagged constructs, and the media was supplemented with 50 uM biotin for 18 hrs.

### Preparation of cell extracts and affinity purification

For standard Western blot (WB) and AP/MS experiments, whole cell extracts were prepared by scraping cells into ice-cold RIPA buffer (50 mM Tris pH 7.5, 150 mM NaCl, 1% NP-40, 0.5% deoxycholate, protease inhibitors), sonicating and clearing by centrifuging at 2800g for 10 min at 4 °C. For BioID experiments, whole cell extracts were prepared in a similar fashion, but the salt concentration in the RIPA was increased to 500 mM.

To increase interactome coverage in the MYPT1 and SPECC1L-GFP AP/MS experiments, separate cytoplasmic and nuclear fractions were prepared for the AP/MS steps and the MS results concatenated at the analysis step [61]. All solutions used in the fractionation, extraction and AP protocols are supplemented with EDTA-free COMPLETE protease inhibitors (Millipore Sigma) to minimize protein degradation. Total protein concentrations for all extracts were measured using a Bradford Assay.For capture of endogenous MYPT1 from cell extracts, MYPT1 antibody (Bethyl Lab) was covalently conjugated to protein A Dynabeads (Invitrogen). Covalently-conjugated affinity purified IgG from the same species (Rabbit; Thermo-Fisher) was used for the control AP. For capture of GFP-tagged proteins from cell extracts, an equal amount of total protein extract for each condition was incubated with GFP-Trap_A beads (Chromotek) at 4°C for 1 hour. Post-AP, the affinity matrices were washed 3x with RIPA buffer and bound proteins eluted with a bead equivalent volume of 1% SDS.

For the BioID experiments, the salt concentration in the lysates was first reduced to 250 mM by adding an equal volume of RIPA buffer with 0 mM NaCl. Equal amounts of total protein extract for each condition were incubated with Streptavidin-agarose beads (ThermoFisher) at 4°C for 4 hours and bound proteins eluted with 2% SDS/30 mM biotin.

For immunoblotting, LDS sample buffer (Invitrogen) was added to lysates and bead eluents and the proteins resolved on a 4–12% Novex Nu-PAGE bis-Tris polyacrylamide gel (Thermo-Fisher) and transferred to nitrocellulose membranes. For the SILAC AP/MS and SILAC BioID/MS experiments, beads from the different conditions were combined pior to the elution step, to minimize variability in downstream processing. The eluted proteins were then reduced and alkylated by treatment with DTT and iodoacetamide, respectively. Sample buffer was added and the proteins resolved by electrophoresis on a NuPAGE 10% BisTris gel (Thermo Fisher). The gel was stained using SimplyBlue Safestain (Thermo Fisher) and the entire lane cut into five slices. Each slice was cut into 2 × 2 mm fragments, destained, and digested overnight at 30°C with Trypsin Gold (ThermoFisher).

### Mass spectrometry and data analysis

An aliquot of each tryptic digest was analyzed by LC-MS/MS on an Orbitrap Fusion Lumos system (Thermo Scientific) coupled to a Dionex UltiMate 3000 RSLC nano HPL. The raw files were searched against the Human UniProt Database using MaxQuant software v1.5.5.1 (http:/www.maxquant.org) [62] and the following criteria: peptide tolerance = 10 ppm, trypsin as the enzyme (two missed cleavages allowed), and carboxyamidomethylation of cysteine as a fixed modification. Variable modifications are oxidation of methionine and N-terminal acetylation. Heavy SILAC labels were Arg10 (R10) and Lys8 (K8). Quantitation of SILAC ratios was based on razor and unique peptides, and the minimum ratio count was 2. The peptide and protein FDR was 0.01. The AP/MS and BioID datasets (minus common environmental contaminants as per http://maxquant.org and proteins identified via the decoy database) are provided in the Supplemental Data Files.

### Fluorescence microscopy

Cells seeded on No. 1.5 coverslips were fixed for 10 min at 37°C in 3.7% (wt/vol) paraformaldehyde (PFA) in PHEM buffer (60mM PIPES pH 6.8, 27 mM HEPES, 20mM EGTA, 16mM MgSO_4_, pH 7.0 with 10M KOH). Following a 10 min permeabilization with 1% Triton X-100 in PBS, nonspecific binding sites were blocked by incubation in PBS with 0.2% Tween-20 and either 1% donkey serum (for immunostaining) or 1% BSA (for Streptavidin staining of biotinylated proteins). For immunostaining, cells were incubated with the appropriate primary antibody, followed by incubation with the appropriate fluorophore-conjugated secondary antibody. Coverslips were prepared for imaging by mounting in Vectashield liquid mounting media (Vector Labs). For BioID, cells were incubated with fluorophore-tagged Streptavidin (Thermo-Fisher).

For live imaging, cells were cultured in 35 um optically clear polymer-bottom u-dishes (ibidi) and growth medium replaced with Phenol Red-free CO_2_ independent medium (ThermoFisher). If desired, DNA was stained by incubating the cells for 20 min at 37°C in medium containing 0.25 μg/ml Hoechst No. 33342 (Millipore Sigma). Images were acquired using a DeltaVision CORE widefield fluorescence system fitted with a 60× NA 1.4 PlanApochromat objective (Olympus), CoolSNAP charge-coupled device (CCD) camera (Roper Scientific) and environmental chamber. The microscope was controlled and images processed by SoftWorX acquisition and deconvolution software (GE Healthcare). All images are single, deconvolved optical sections.

Photobleaching experiments were carried out as previously described [63]. Briefly, cells were cultured in glass-bottom dishes (Ted Pella). Three single sections were imaged before photobleaching, a region of interest (ROI) then bleached to ∼50% of its original intensity using the 488-nm laser, and a rapid series of images acquired after the photobleaching period for a total of 90 sec. Recovery curves were plotted and the mobile fractions and half times of recovery determined using GraphPad Prism.

### Super-resolution imaging

Expansion Microscopy (ExM)-based super-resolution imaging was carried out as per our modification [43] of the X10 protocol [42]. Briefly, cells seeded on coverslips were transfected for 18 hrs to drive transient overexpression of SPECC1L-GFP. They were then PFA-fixed and permeabilized, and stained with anti-acetylated-alpha-tubulin and anti-mouse-Alexa647. To counteract the reduction in GFP signal following expansion, cells were also stained with rabbit anti-GFP and anti-rabbit-Alexa488 antibodies. The stained cells were then incubated overnight incubation at room temperature with Acryloyl-X anchoring reagent (ThermoFisher). The gelation solution was prepared by mixing 1.335g DMAA and 0.32 g sodium acrylate (Millipore Sigma) with 2.85 g ddH_2_O, vortexing and purging O_2_ by bubbling with N_2_ for 40 min at room temperature. A solution of 0.036 g/ml of potassium persulfate was prepared in ddH2O, 0.3 ml added to 2.7 ml of gelation solution and O_2_ purged by bubbling with N_2_ for 15 min on ice. Gelation chambers were prepared by sandwiching the stained coverslip, cell side up, between two coverslips and adding spacer coverslips along the sides. TEMED was added to the gelation mix, 100 ul of the solution pipetted on top of the cells using a pre-chilled pipette tip and a 22 × 22 mm coverslip placed on top. Gels were placed in a humidified chamber and allowed to polymerize for 3 hours at room temperature. The gels were removed from the coverslips incubated overnight at 50℃ with 8 U/ml Proteinase K (Sigma-Aldrich) prepared fresh in digestion buffer (50mM TRIS, 800mM Guanidine HCl, 1mM EDTA and 0.5% (v/v) Triton X-100 in ddH2O, pH 8.0). To swell the gels, deionized water containing 1.7 ug/ml Hoechst 33342 was added. After 10 min, this was replaced with 2 more washes of deionized water for a total of 1 hour expansion. To image cells post-expansion, segments were removed using a custom 3D-printer gel cutter [43] and transferred to 8-well chambered coverslips (ibidi). Multiwavelength Z-stacks were acquired using a Zeiss LSM880 confocal laser scanning microscope with a 63x/1.4 NA Plan-Apo objective in AiryScan mode.

For dSTORM super-resolution imaging, cells seeded on No 1.5 coverslips were transfected for 18 hrs to drive transient over expression of Specc1L-GFP, and then PFA-fixed, permeabilized and stained with anti-acetylated-alpha-tubulin and anti-mouse-Alexa647. The coverslips were mounted in a 40% Vectashield, 60% Tris/glycerol solution that has been shown to support efficient blinking of Alexa647 [64]. Images were acquired on a Quorum spinning disk confocal system (Quorum Technologies) using a 63X/1.4NA objective, a Hamamatsu EM-CCD ImagEM camera and the appropriate laser lines and filter sets. Initially, widefield and spinning disk confocal images were acquired for both the SPECC1L-GFP signal (490 nm laser and FITC emission filter set) and the anti-acetylated-alpha-tubulin-Alexa647 signal (639 nm laser and Cy5 emission filter set) using low laser power. A region of interest was then chosen for super-resolution imaging, using the MetaMorph Single Molecule Resolution real time acquisition and analysis module (Molecular Devices LLC). Stochastic activation of the Alexa647 fluorophores was achieved by firing the 639 nm laser at full power, which causes them to blink as they cycled through their ON/OFF states. A series of 10,000 images was acquired, and a super-resolved image reconstructed from the individual x-y localizations mapped in each image.

### Purification of recombinant proteins for in vitro assays

Plasmids expressing recombinant 6x His-tagged SPECC1L fragments were transformed in BL-21 cells and plated on LB/ampicillin plates overnight at 37°C. LB was inoculated with a colony and expanded to a larger culture that was induced with 0.5 mM IPTG once the OD_600_ reached ∼0.5. Following 18 hrs incubation at 16°C, the culture was pelleted, resuspended in buffer (20 mM Tris pH 7.5, 250 mM NaCl, 10 mM Imidazole) containing 1 mg/mL of Lysozyme and incubated on ice for 30 min. Following sonication and pelleting of cell debris, the lysate was incubated with Ni^2+^ -NTA agarose beads (Qiage) at 4°C for 1 hr with end-over-end rotation. The beads were pelleted and washed 3x with wash buffer (20 mM Tris pH7.5, 0.5 M NaCl, and 20 mM Imidazole). The His-tagged proteins were eluted with elution buffer (20 mM Tris pH7.5, 0.5 M NaCl, and 300 mM Imidazole) at 500 μl increments and dialyzed in TGEM buffer (20 mM Tris-HCl pH7.9, 0.1 M NaCl, 20% glycerol, 1mM EDTA, 5 mM MgCl_2_, 0.1% NP-40, 0.2 mM PMSF). After dialysis 1 mM DTT was added prior to use in the in vitro assays.

Plasmids expressing recombinant GST-tagged Mypt1(CD) or GST alone were transformed in Rosetta™ cells and plated on LB/ampicillin plates overnight at 37°C. LB was inoculated with a colony and expanded to larger culture that was induced with 0.2 mM-0.5 mM IPTG once the OD_600_ reached ∼0.5. Following 18 hrs incubation at 16°C, the culture was pelleted and resuspended in buffer (50 mM Tris pH 8.0, 2 mM EDTA pH 8.0, 0.1% BME, and protease inhibitor). Following sonication and pelleting of cell debris, NaCl was added to a final concentration of 0.25 M and the lysate was incubated with Glutathione sepharose beads (GE Healthcare) at 4°C for 1 hr with end-over-end rotation. The beads were pelleted and washed 3x with wash buffer (20 mM Tris pH7.5, 0.25 M NaCl, 2 mM EDTA, 2 mM EGTA). GST-Mypt1(CD) was eluted using 10 ml wash buffer containing 61 mg of reduced glutathione and 13 μl of 10 M NaOH (pH 8.0). The eluted protein was dialyzed in TGEM buffer (20 mM Tris-HCl pH 7.9, 0.1 M NaCl, 20% glycerol, 1 mM EDTA, 5 mM MgCl_2_, 0.1% NP-40, 0.2 mM PMSF). After dialysis 1 mM DTT was added prior to use in the in vitro assays.

### In vitro co-immunoprecipitation assay

Purified recombinant HisSpecc1LΔCD was combined with either GST and GST-Mypt1(CD) immobilized on Glutathione agarose beads. Following 1 hr incubation at 4C, bound protein was eluted, subjected to SDS-PAGE and transferred to nitrocellulose membrane for immunoblotting with anti-His and anti-mouse-HRP to detect the Specc1L fragment. The blot was then stripped and probed with anti-GST and anti-mouse-HRP.

### Far Western blot assay

Purified recombinant HisSpecc1LΔCD (6 ug) was separated by SDS-PAGE and transferred to nitrocellulose membrane. The membrane was incubated overnight with 10% milk (in PBS + 0.5 % Tween) to block nonspecific binding sites and washed 3x with PBS/0.5%Tween, then 1x with PBS, before it was overlaid with 5 μg purified recombinant GST-Mypt (CD) or GST in 5 mL of 1 mg/mL BSA/PBS solution. The blot was then probed with anti-GST and anti-mouse HRP.

### Microtubule binding protein spin-down assay

The Microtubule Binding Protein Spin-down assay was performed according to the manufacturers’ protocol (Cytoskeleton). Briefly, stabilized microtubules were prepared by incubating a 20 ul aliquot of Tubulin protein (5 mg/mL) at 35°C with 2 ul of Cushion Buffer (80 mM PIPES pH7.0, 1 mM MgCl2, 1 mM EGTA, 60 % glycerol) for 20 min on ice. 2 ul of 2 mM Taxol was added to 200 ul of preheated (at 35°C) General Tubulin Buffer (80 mM PIPES pH 7.0, 2 mM MgCl2, 0.5 mM EGTA), which was immediately added to the MTs after the 20 min incubation on ice. The sample was gently and thoroughly mixed and left at room temperature, allowing the MTs to stabilize. 20 ul of the stable microtubules were then mixed with either Bovine serum albumin (BSA), MAP4, or His-SPECC1L-NT (2-5 ug). After a 30min incubation the samples were carefully pipetted onto 100 ul of Cushion Buffer (1 mL containing 10 ul of 2 mM Taxol) and centrifuged at 100, 000 x g at room temperature for 40 min. The uppermost supernatant layer (50 ul) was removed and added to 16.7 ul of 4x Laemmli sample buffer. The remaining supernatant was discarded. The pellet was resuspended in 66.7 ul of 1x Laemmli sample Buffer. 20 ul of the supernatant and pellet sample were subjected to SDS-PAGE and either Coomassie stained or transferred to nitrocellulose for immunoblotting.

### Actin Fractionation Assay

U2OS cells co-transfected with Flag-tagged Actin (wild type or R62D nonpolymerizable mutant) and GFP-tagged constructs were washed in PBS and extracted with 0. 5 ml of cytoskeletal buffer (50 mM MES pH 6.8, 1 mM EGTA, 50 mM KCl, 1 mM MgCl2, 0.5% Triton X-100, protease inhibitors) on a shaking incubator. After 2 min, the Triton X-100-soluble fraction was removed and the Triton X-100 insoluble fraction was scraped off the dishes into 0.5 ml of the same buffer and sonicated. Aliquots of each fraction (100 ul) were mixed with equivalent amount of LDS sample buffer and reduced to a final volume of 45 ul in a SpeedVac for separation by 10% SDS-PAGE, transfer to nitrocellulose and immunoblot analysis using anti-Flag and anti-GFP antibodies.

## ACKNOWLEDGMENTS

We thank colleagues in the Trinkle-Mulcahy lab for helpful discussions and suggestions, and Priyam Maini and Sarah Ooi for their technical contributions. We also thank Drs. Ina Poser and Anthony Hyman for reagents, and Lawrence Puente at the Ottawa Hospital Research Institute Proteomics Core Facility for technical support. We acknowledge the outstanding support of the Cell Biology and Image Acquisition Core (RRID: SCR_021845) funded by the University of Ottawa and the Canada Foundation for Innovation. Funding: This work was supported by Natural Sciences and Engineering Research Council Discovery Grant 06674 (to L.T.-M.) and a Canadian Institutes of Health Research Banting & Best Scholarship (to V.M).

## AUTHOR CONTRIBUTIONS

V.M. and L.T.-M. planned the experiments. V.M., A.G.L. and N.D. performed the experiments. V.M., J.C., A.G.L. and L.T.-M. contributed intellectually to the work as well as read and edited the final manuscript.

### COMPETING INTERESTS

The authors declare that they have no competing interests.

## DATA AND MATERIALS AVAILABILITY

All data needed to evaluate the conclusions in the paper are present in the paper or the Supplementary Materials. Plasmids and cell lines are available through Addgene or upon request.

## SUPPLEMENTAL FIGURES and TABLE

**Figure S1.**
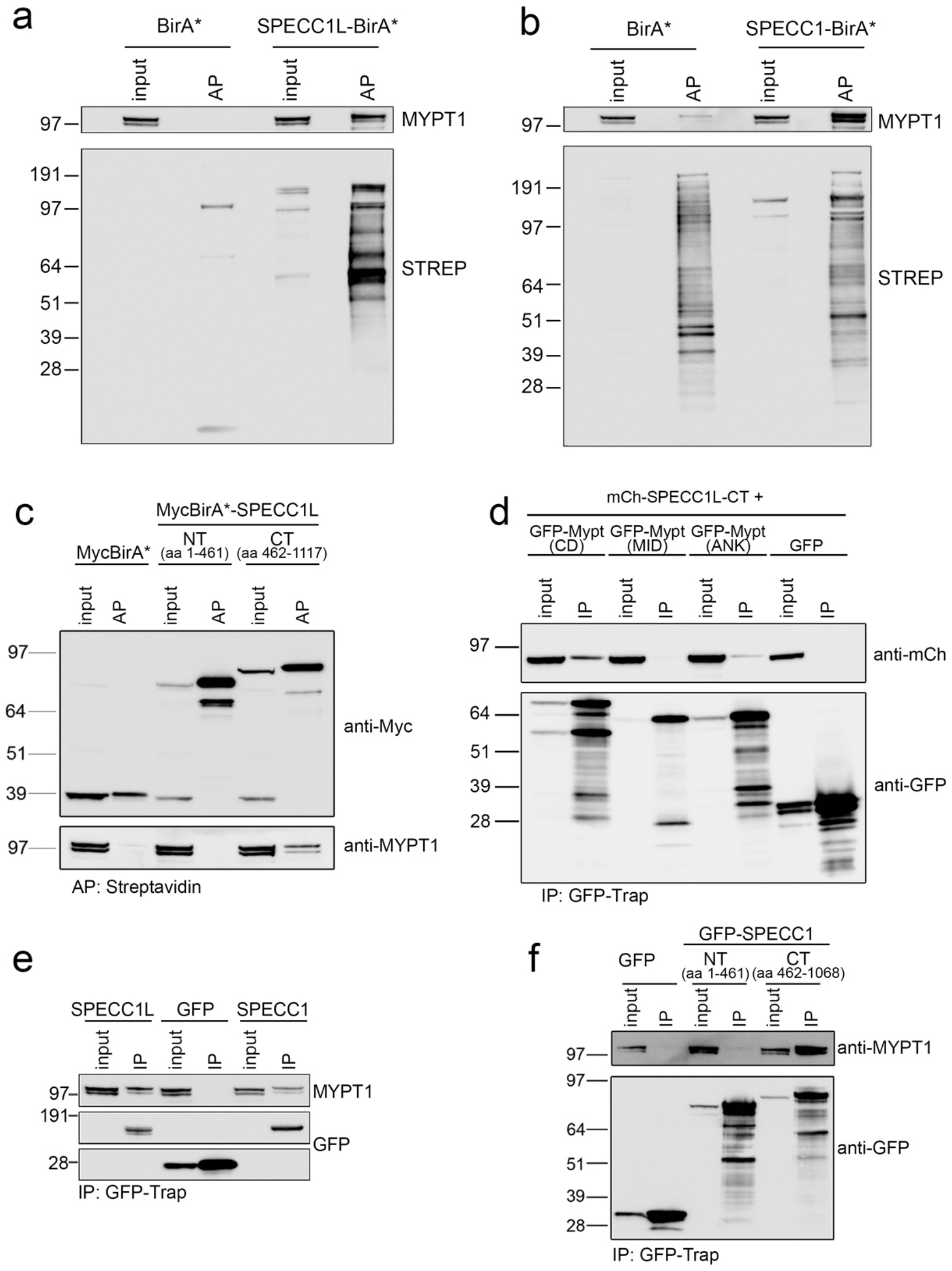
Validation of the interaction of MYPT1 with SPECC1L and SPECC1. a. Proximity biotin labeling of endogenous MYPT1 by Myc-BirA*-SPECC1L and Myc-BirA*-SPECC1 expressed in U2OS cells allows it to be captured on Streptavidin-agarose beads and visualized by WB analysis (a-b). Top panels were probed with anti-MYPT1 and bottom panels with Streptavidin-HRP. c. Only the C-terminal half of SPECC1 is able to proximity label MYPT1 when expressed as a Myc-BirA* fusion protein. The top panel was probed with anti-Myc and the bottom panel with anti-MYPT1. d. When co-expressed in cells, the GFP-MYPT1 (CD) fragment co-immunoprecipitates mCh-SPECC1L-CT. e. GFP-tagged SPECC1L and SPECC1 both co-purify endogenous MYPT1 from U2OS cell lysates. f. Only the C-terminal half of SPECC1 co-immunoprecipitates endogenous MYPT1. These data are related to Figures 1-2.

**Figure S2.**
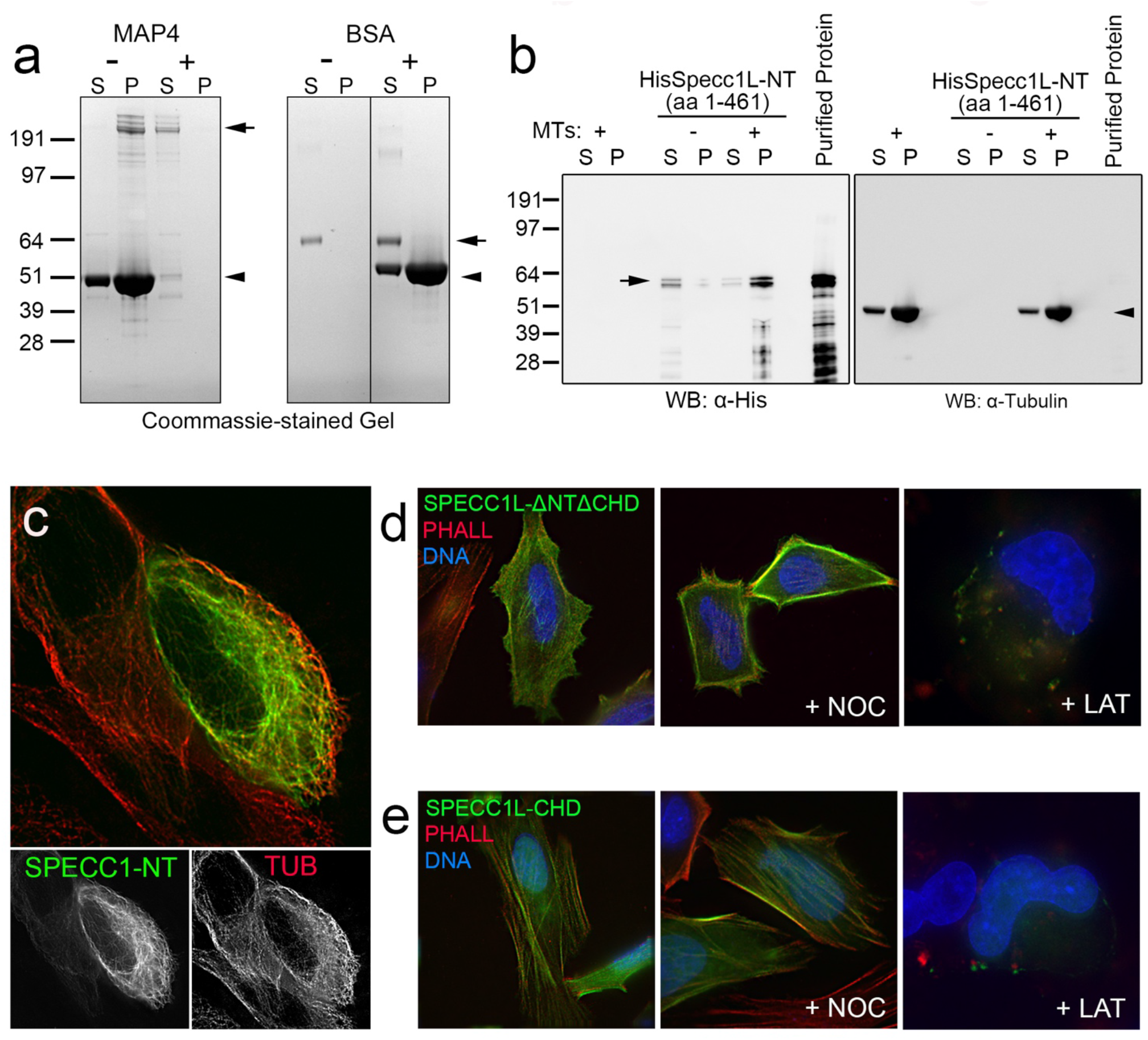
MT binding assays, SPECC1-NT MT localization and sensitivity of SPECC1L C-terminal fragments to LAT treatment. a. Positive (MAP4) and negative (BSA) controls used for the microtubule binding assay shown in Fig. 3. b. WB blot run in parallel to the His-SPECC1L-NT MT binding assay Coomassie-stained gel and probed with anti-His and anti-alpha-tubulin to confirm that the stained bands represent the recombinant protein and tubulin, respectively. c. The N-terminal half of SPECC1 (SPECC1-NT, aa 1-461; green) governs its association with MTs (red). Treatment of the SPECC1L-deltaΔNTΔCHD (aa 462-890, d) and SPECC1L-CHD (aa 891-1117, e) truncation mutants confirms that their localization patterns (green), which co-localize with Phalloidin-stained actin filaments (red), are sensitive to Latrunculin B (LAT) but not Nocodazole (NOC) treatment. Hoechst 33342-stained DNA is shown in blue. These data are related to Figure 3. Scale bars are 10 μm.

**Figure S3.**
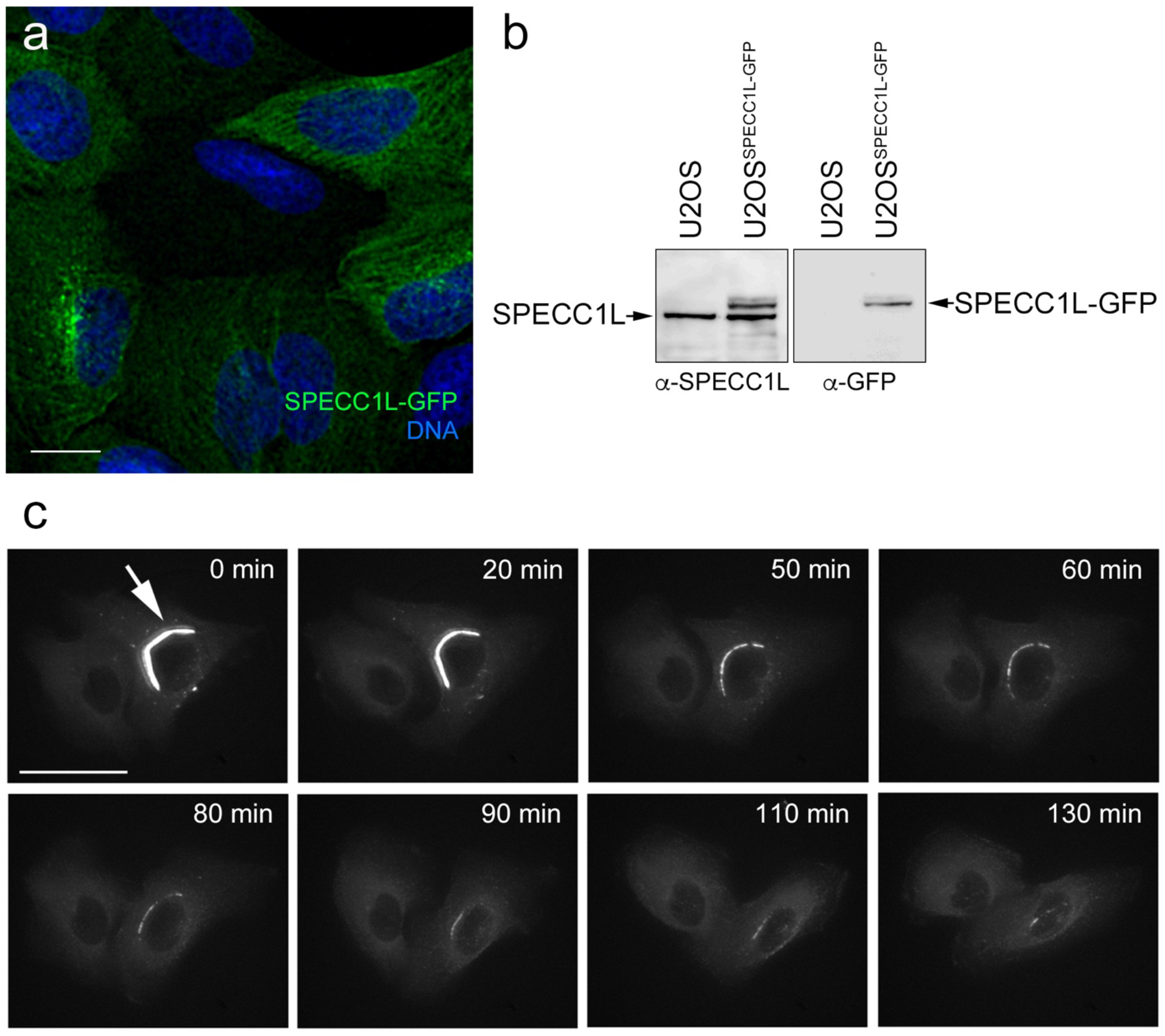
Characterization of U2OS^SPECC1L-GFP^ stable cell line. a. Field of view of U2OS cells stably overexpressing SPECC1L-GFP (green). Hoechst 33342-stained DNA is shown in blue. Scale bar is 10 μm. b. WB analysis with anti-GFP and anti-SPECC1L antibodies shows that the U2OS^SPECC1L-GFP^ stable line expresses the fusion protein at a similar level to the endogenous protein. c. Live imaging shows that bundled MTs (arrow) are occasionally observed in U2OS^SPECC1L-GFP^ cells but resolve over time. Scale bar is 15 μm. These data are related to Figure 4.

**Figure S4.**
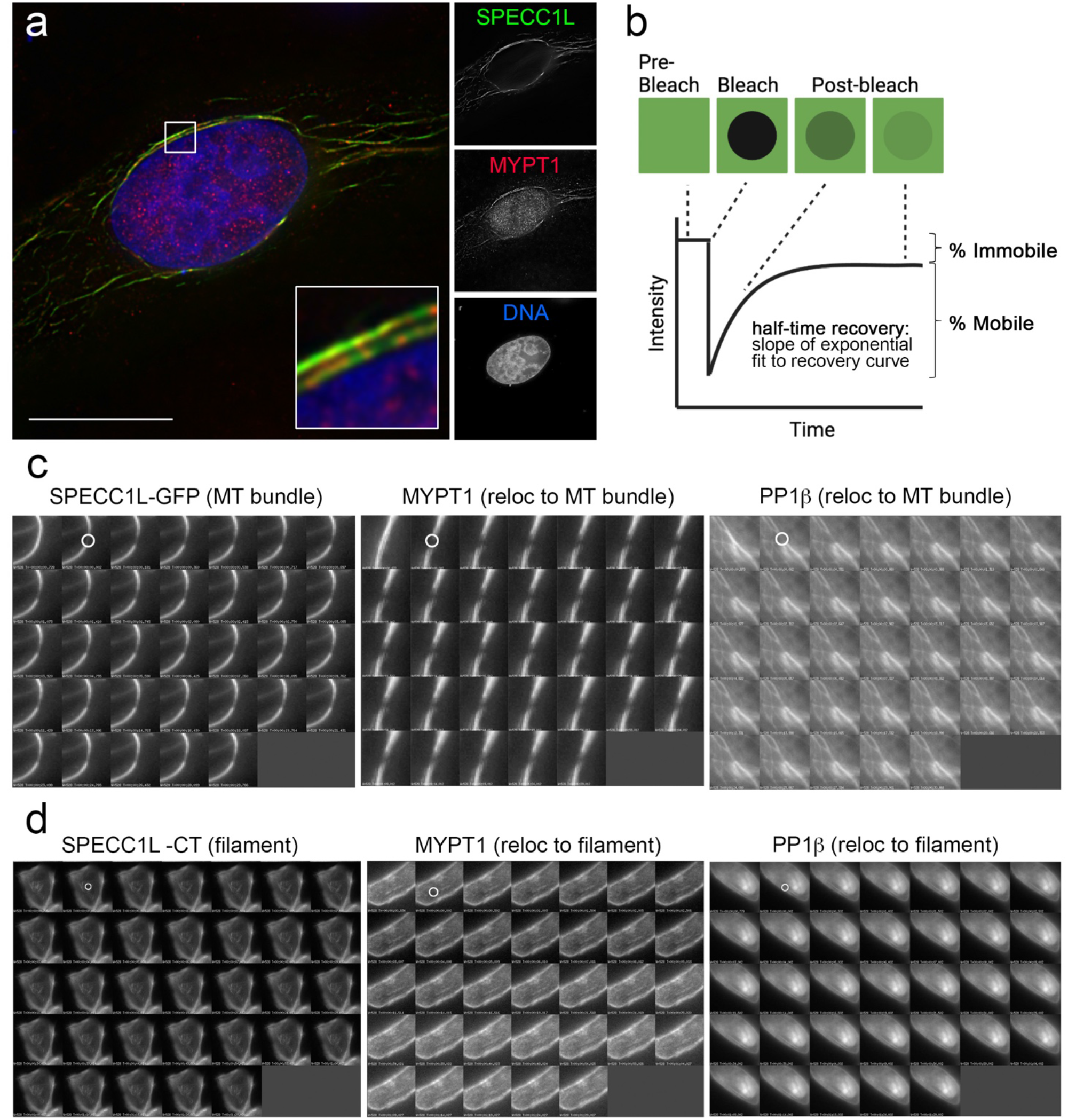
SPECC1L modulates the subcellular distribution and turnover dynamics of the myosin phosphatase complex. a. Endogenous MYPT1 (red) accumulates at the bundled/stabilized MTs induced by transient overexpression of SPECC1L-GFP (green). Scale bar is 10 μm. b. Diagram summarizing the design of the FRAP experiments. Following a pre-bleach image, the GFP in a region of interest (ROI) is photobleached at 100% laser power and recovery of signal in the ROI monitored over time (90 sec post-bleach). c. Examples of FRAP experiments for SPECC1L-GFP accumulated at a MT bundle, and Mypt1-GFP and PP1β-GFP relocalized to MT bundles by SPECC1L. d. Examples of FRAP experiments for GFP-tagged SPECC1L-CT accumulated at actin filaments, and Mypt1-GFP and PP1 β -GFP relocalized to filament structures by SPECC1L-CT. In each panel the ROI is indicated by a white circle in the first post-bleach image. These data are related to Figure 5.

**Figure S5.**
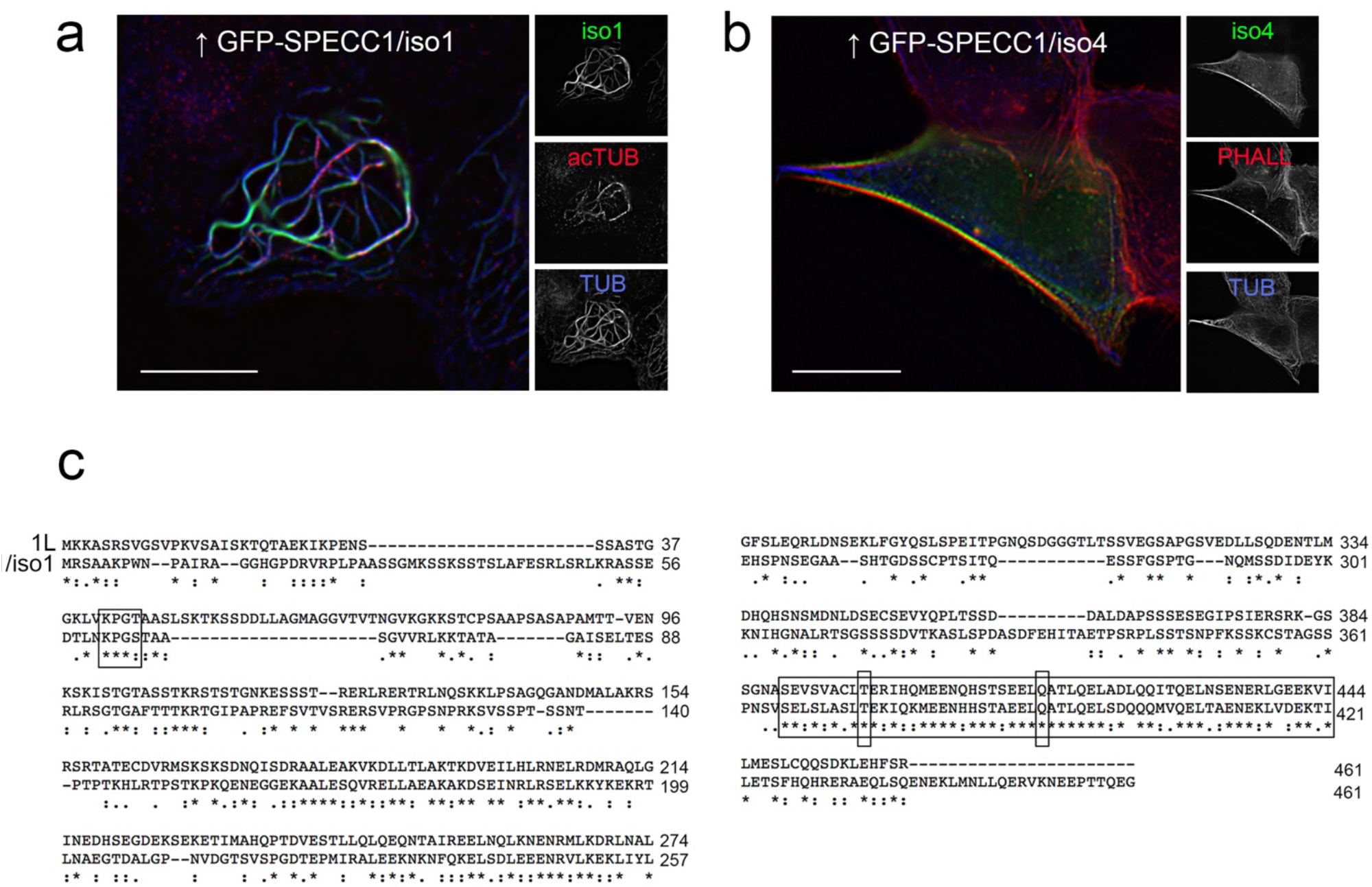
Putative microtubule binding regions in SPECC1L and SPECC1. SPECC1/iso 1 (green) induces MT stabilization/bundling (anti-Tub in blue, anti-acTub in red) when transiently overexpressed in cells (a), while the splice variant SPECC11/iso4 (b) remains associated with actin filaments (phalloidin, red). Scale bars are 10 μm c. Clustal Omega alignment of aa 1-461 in SPECC1L and SPECC1/iso1. The large box highlights the region of similarity (>70% identical) in CCD2. Missense mutations within this region have been identified in patients with developmental disorders (T396P and Q415P indicated here). The putative MT-binding KXGS/T motif that lies closer to the N-terminus of both SPECC1L and SPECC1/iso1 (and is not found in SPECC1/iso4) is indicated by the smaller box. These data are related to Figure 4.

**Figure S6.**
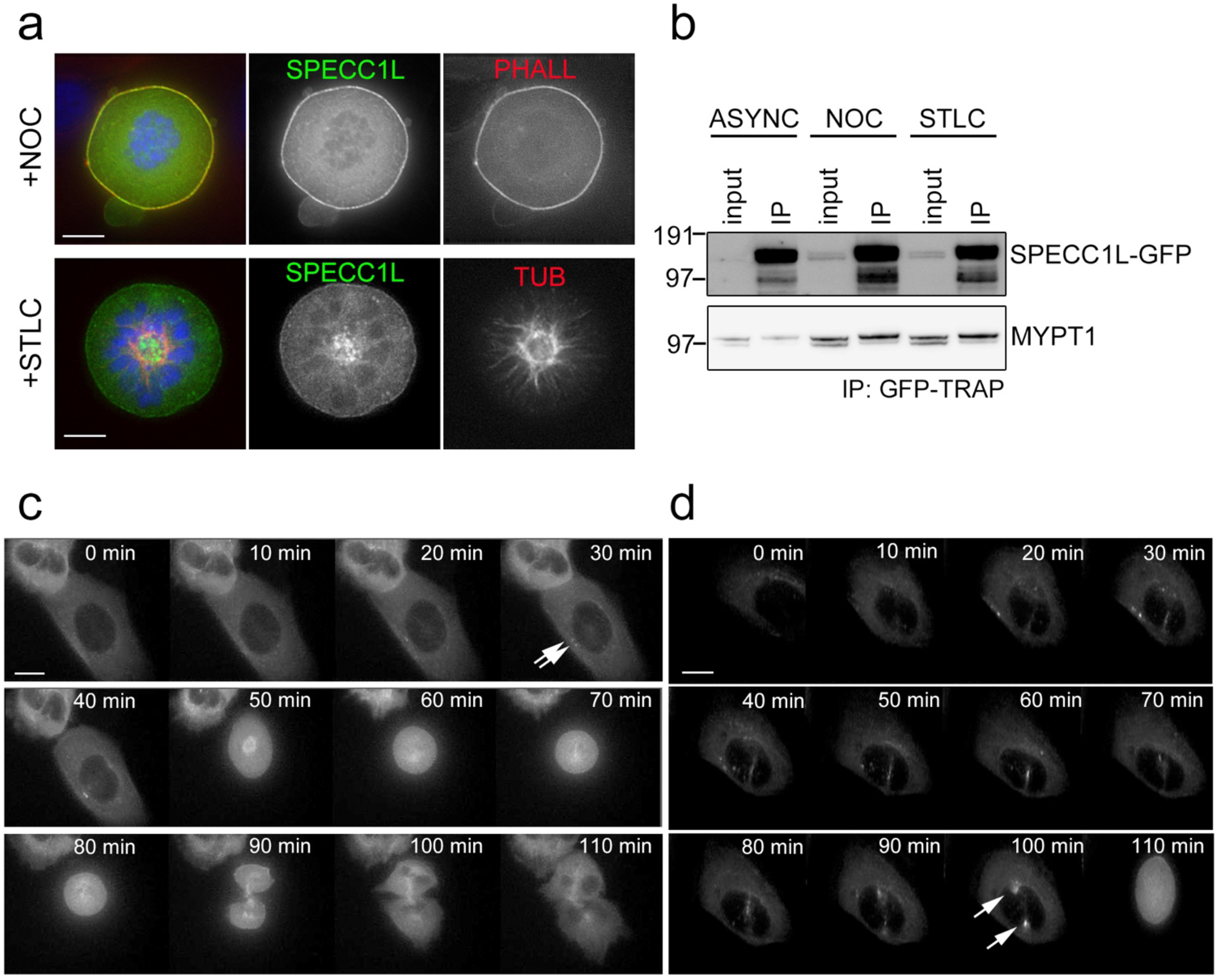
SPECC1L associates with MT and actin structures, and with MYPT1, throughout the cell cycle. a. In NOC-arrested U2OS^SPECC1L-GFP^ cells, SPECC1L-GFP (green) is diffuse in the cytoplasm, remaining distinct from condensed chromosomes (blue) and showing additional accumulation with filamentous actin (Phalloidin, red) at the cell cortex. In STLC-arrested U2OS^SPECC1L-GFP^ cells, SPECC1L-GFP (green) is diffuse in the cytoplasm, remaining distinct from condensed chromosomes (blue) and accumulating at the spindle pole region at the center of the anti-tubulin stained monoastral spindle (red). b. IP/WB analysis confirms association of MYPT1 with SPECC1L throughout the cell cycle. Panels c-d show examples of the transient association of SPECC1L-GFP with centrosomes (arrows) at the onset of mitosis in live imaging experiments. Scale bars are 10 μm These data are related to Figure 6.

**Table S1.** Mobile fraction and half-time recovery values calculated for GFP-tagged SPECC1L, Mypt1 and PP1β. GraphPad Prism was used to fit double exponential curves to the 90 sec post-bleach time courses shown in Fig. 5c. The immobile fraction was determined (% that does not recover within 90 sec), and the distribution and recovery rates for the two mobile fractions (slow and fast) characterized. The R2 value for all curve fits was >0.95.

| CONDITION | Immobile Fraction (%) | SLOW POOL |  | FAST POOL |  |
| --- | --- | --- | --- | --- | --- |
|  |  | % Recovered | Half-time (sec) | % Recovered | Half-time (sec) |
| Free GFP |  |  |  |  |  |
| Unperturbed | 0.9 | 8.9 | 1.14 | 90.2 | 0.00063 |
| SPECC1L-GFP |  |  |  |  |  |
| Unperturbed | 3.7 | 19.9 | 1.67 | 76.4 | 0.00072 |
| MT bundles (↑ SPECC1L) | 54.7 | 17.8 | 21.78 | 27.5 | 0.00092 |
| Filaments (↑ SPECC1L-CT) | 17.5 | 27.9 | 20.69 | 54.6 | 0.00056 |
| Mypt1-GFP |  |  |  |  |  |
| Unperturbed | 8.4 | 19.2 | 4.23 | 72.4 | 0.00065 |
| MT bundles (↑ SPECC1L) | 57 | 6.9 | 25.32 | 36.1 | 0.16 |
| Filaments (↑ SPECC1L-CT) | 37.1 | 9.4 | 9.96 | 53.5 | 0.00058 |
| PP1β-GFP |  |  |  |  |  |
| Unperturbed | 2.1 | 16.3 | 2.8 | 81.6 | 0.00008 |
| MT bundles (↑ SPECC1L) | 26.7 | 1.73 | 9.42 | 56 | 0.00092 |
| Filaments (↑ SPECC1L-CT) | 17.2 | 18.3 | 12.6 | 6.43 | 0.0008 |

## Notes

### Competing Interest Statement

The authors have declared no competing interest.

